# Interspecies interaction controls *Escherichia coli* growth in human gut microbiome samples

**DOI:** 10.1101/2024.09.27.615362

**Authors:** Mathilde Boumasmoud, Ricardo León-Sampedro, Vera Beusch, Fabienne Benz, Markus Arnoldini, Alex R. Hall

## Abstract

Gut microbial community composition varies from one person to another. Potentially, this means the ecological interactions experienced by individual strains or species also vary among microbiomes of different people. However, testing this directly in human microbiomes and identifying ecological drivers involved is challenging. Here, we use replicated anaerobic microcosms to quantify variability of population growth for a key commensal species among microbiome samples from different individuals, and to identify underlying intra- and interspecific interactions. In a reciprocal transplant experiment, both absolute and relative growth performance of different *Escherichia coli* strains varied among gut microbiome samples from healthy individuals. This was partly explained by intraspecific competition: ecological success of individual *E. coli* strains was associated with displacement of resident conspecifics. However, the determinants of *E. coli* growth varied among samples. In one microbiome sample with a distinctive taxonomic composition, culture acidification by resident microbes impaired growth of all *E. coli* strains. We identified a strain of *Clostridium butyricum* contributing to this effect, and showed that transferring it into other microbiomes predictably altered pH, fermentation product profiles (butyrate accumulation and acetate/lactate depletion) and population growth of other species including *E. coli*, thereby reshaping overall taxonomic composition. Our results suggest natural inter-individual gut microbiome variation translates to variable ecological interactions with incoming bacteria, but these dynamics can be manipulated by a generalizable interspecies interaction.

**Significance statement:** Gut-microbiome variation among healthy individuals is widely documented, yet its impact on the success of incoming bacterial strains and its implications for microbiome-targeted interventions remain unclear. Here, we test this experimentally by cultivating stool samples from healthy individuals and transplanting strains across samples. We reveal a functional relationship between microbiome variation and growth performance of individual strains. This underscores the challenge of predicting colonization outcomes. Yet, we identify a taxon contributing to the distinct functional profile of one of the samples, and show that its transplantion into other microbiome samples reproducibly alters key properties, including pH and population growth of other bacteria, revealing a common ecological control point. These findings advance our ability to explain and manipulate human gut-microbiome variation.

## Introduction

Microbial colonization of the human gastrointestinal tract begins at birth [1]. After a dynamic development during the first years of life [2], microbial community composition remains relatively stable, with changes mostly in the relative abundances of different taxa [3]. Within individual subjects, resident strains can be retained for extended periods [4,5] but also occasionally replaced by newly-arriving strains [6,7]. Among individual subjects, taxonomic composition of the gut microbiota is highly individual, as evidenced by community profiles from 16S rRNA amplicon sequencing and analyses at the strain-level [8]. This variation among microbiomes of different individuals, shaped by host and environmental factors [2,9], may itself be a key determinant of who becomes colonized by which newly incoming strain, both early on in life and during later strain turnover. In support of this idea, various mechanisms have been identified by which interactions with resident microbiota can affect population growth of individual strains, such as nutrient competition [10], crossfeeding [11] and direct inhibition [12]. Thus, variable taxonomic composition may result in variation of these ecological interactions and, in turn, variable population growth of incoming strains. Experiments in mice support this idea: colonization success for the same strain can vary across different microbiomes [13]. Furthermore, different strains within a given species often have different phenotypes relevant for colonisation, such as nutrient preferences [14]. Some of these strain-specific traits may be advantageous in some microbiomes but not in others. This means that inter-individual microbiome variation may determine both the relative performance of different strains within species, and average population growth at the species level.

Here, we quantify variation in population growth of the key gut commensal *Escherichia coli* across gut microbiome samples from different healthy individuals, including a resident microbiota and abiotic components [15]. First, we focus on the strain level to ask how absolute and relative performance of different strains varies among microbiome samples. Quantifying this variability is important for understanding the ecological determinants of strain diversity. For example, if relative performance of different strains varies among samples from different people, then microbiome-to-microbiome variation may be a key determinant of colonisation success for incoming strains and subsequent turnover. Furthermore, if the identity of the best-performing strain varies among microbiome samples, a pattern of local adaptation could emerge [16,17]. Despite many tests in other organisms, including plants and animals [18–20], and microbes in other ecosystems [21–25], it remains unclear whether such patterns apply to bacteria in human microbiomes. Second, we focus on the species level and ask what type of bacterial interactions explain variable average population growth of *E. coli* among microbiome samples from different individuals. Like many other species, *E. coli* naturally makes up a variable proportion of resident microbiota [26], raising the question of whether its abundance in a given microbiome is strongly influenced by interactions with particular taxa [27]. This aspect is relevant in the context of potential microbiome-based therapies, such as probiotics and fecal microbiota transplantation [28,29]. For instance, if growth of a focal species is strongly affected by interactions with one or a few other taxa, its abundance could potentially be manipulated therapeutically by inhibiting or introducing such relevant taxa [30]. By contrast, effects arising from complex interactions among several strains or species would be harder to manipulate predictably, requiring a more personalised approach instead [31].

Addressing these questions directly in human microbiomes and identifying the underlying ecological drivers is challenging. There have been major advances on related questions from work with synthetic communities in microcosms or gnotobiotic hosts [32–35]. However, the genetic variation within artificial communities, and differences among experimental groups in such systems, are not reflective of natural human-associated communities or real-world person-to-person variation. For example, the set of strains in an assembled community is typically not as species-rich and does not share the same history of ecological interactions as a natural community. Similarly, experiments with mice and associated microbiota have produced important insights [13,36], but leave gaps in our understanding of human-associated communities [37,38]. An important practical obstacle is that it is extremely challenging to measure the population growth of individual strains in the gut microbiomes of healthy individuals, let alone to do so under replicated introductions of the same strain set into the same microbiomes. We circumvent some of these challenges by using anaerobic microcosms constituted from fecal microbiome samples of healthy humans [39]. This enabled us to perform a reciprocal transplant experiment [16], measuring growth performance of six commensal *E. coli* isolates across the microbiome samples they were isolated from. Thus, we quantify the contribution of strain-to-strain and microbiome-to-microbiome differences to the ‘ecological success’ of commensal *E. coli*, interpreting ecological success or performance as the capacity to reach high abundance in the presence of a resident microbiota. We then investigate the underlying intra- and interspecies interactions and identify a single taxon capable of consistently altering the functional profile of different microbiome samples, thereby controlling growth of other bacteria including *E. coli*.

## Results

### Resident microbiota drive strain-specific *E. coli* performance across microbiome samples from different individuals

We sampled gut microbiomes from the stool of six healthy humans and from each we isolated a single *Escherichia coli* clone, representative of the most abundant resident *E. coli* strain (see Methods, Supplementary Fig. 1). We then inoculated these six focal strains, tagged with a selective marker, into anaerobic microcosms prepared with each of the six microbiome samples resuspended in a basal medium, generating 36 unique focal strain-microbiome combinations. Within 24 h, all focal strains reached high abundance (Fig. 1a). However, their final abundance varied both among focal strains and among microbiome samples from different individuals (F_5,72_=27.4, *p* < 0.001 and F_5,72_=15.0, *p* < 0.001 respectively in a two-way ANOVA, Fig. 1a). Furthermore, the relative performance of different focal strains depended on the microbiome context (interaction term, F_25,72_=2.2, *p*=0.005). In some microbiome samples, such as M3, strain-to-strain differences were large, whereas in others, such as M6, strains performed more similarly. Despite these differences in the magnitude of variation, the rank order of strain performance was generally consistent across microbiome samples, with strains S3 and S7 generally reaching the highest abundances (Fig. 1a). Testing for evidence of local adaptation revealed no average difference between sympatric and allopatric strain-microbiome combinations (Supplementary Fig. 2a). Thus, some focal strains were generally more successful than others in terms of population growth in microcosms with a resident microbiota, but the strength of these differences and average ecological success varied across microbiome samples from different individuals.

**Fig. 1:**
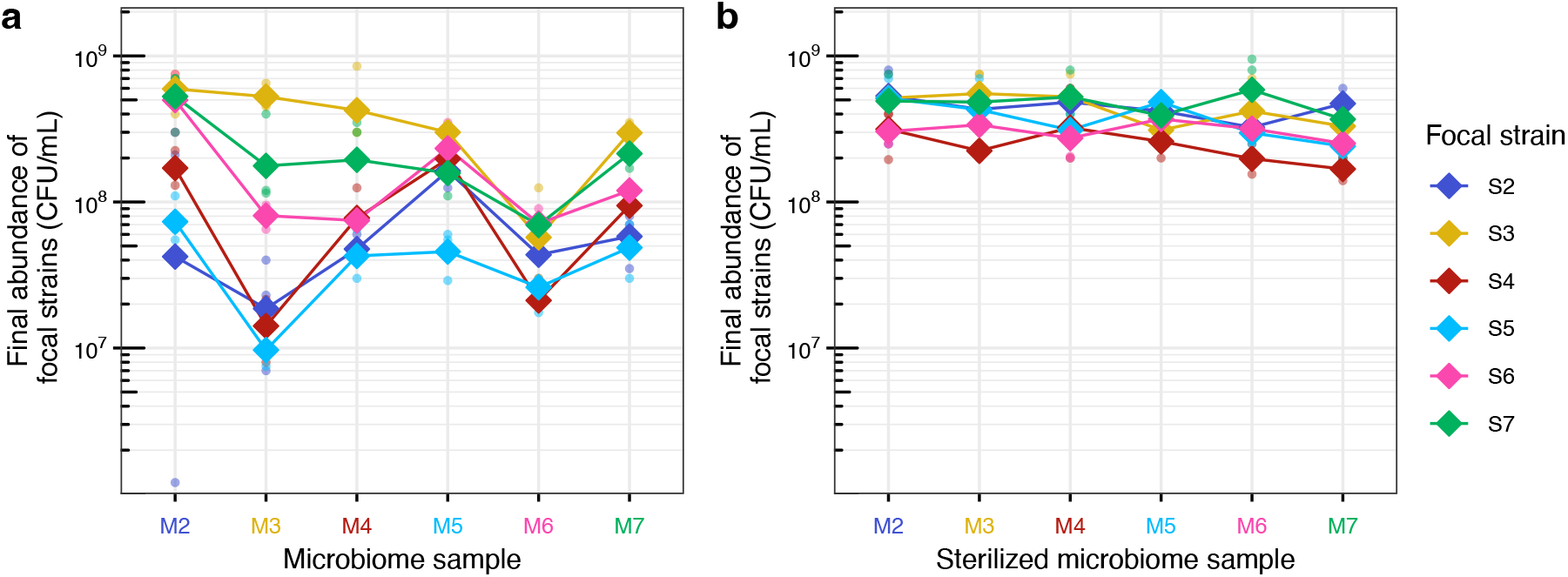
Growth performance of focal *E. coli* strains in human gut microcosms. Final abundance of each focal *E. coli* strain (S2-S7) in microcosms prepared with each microbiome sample (M2-M7) live (a) or sterilized (b), estimated by selective plating after 24h of anaerobic incubation. Diamonds represent the average (geometric mean) of three replicates for each strain-microbiome combination and dots show individual replicates. Lines connect the average for each strain across microbiome samples. Colors of focal strains are matched to the color of their microbiome of origin (e.g., strain 3, S3, was isolated originally from microbiome sample 3, M3).

To test for evidence that variable performance of *E. coli* in microbiome samples was linked to interactions between focal strains and resident microbiota, we performed an experiment equivalent to that above, but with sterilized versions of the same microbiome samples (hereafter, ‘sterile’ microcosms, as opposed to ‘live’ microcosms in the main experiment above). Here, all focal strains reached high abundance in all sterile microcosms (>10^8^ CFU/mL, Fig. 1b). Relative performance of different focal strains was stable among sterilized microbiome samples (interaction term F_25,72_=0.86, p=0.66), and the range of final abundances was smaller than in the experiment above with live microcosms (ranging from 1.7 × 10^8^ to 5.9 × 10^8^ CFU/mL vs. 9.7 × 10^6^ to 5.9 × 10^8^ CFU/mL). We also found no evidence of local adaptation in these conditions (Supplementary Fig. 2b). Nevertheless, we still found average differences among focal strains and sterilized microbiome samples (F_5,72_= 9.2, *p* < 0.001 and F_5,72_=3.1, p=0.012 respectively). There was no overall association between final abundances in sterile microcosms and in live microcosms (R=0.25, p=0.15, Supplementary Fig. 3), suggesting the drivers of focal strain performance were different in the presence versus absence of resident microbiota. In summary, the contrast between the large variability observed in live microcosms (Fig. 1a) and the invariably high abundances across strain-microbiome combinations in sterile microcosms (Fig. 1b) supports a critical role of the resident microbiota in determining strain-specific performance across microbiome samples from different individuals.

### Intraspecific competition shapes strain performance within microbiome-specific finite *E. coli* abundance

We next tested for further evidence of interactions between focal strains and resident microbiota. First, we focused on their interactions with resident *E. coli* in each microbiome sample. Total *E. coli* abundance (including focal strain and resident *E. coli*) in live microcosms varied among samples from different individuals (F_5,72_= 76.7, *p* < 0.001, Fig. 2a). Within each microbiome sample, total *E. coli* abundance was unaffected by focal strain identity (F_5,72_= 1.3, p= 0.25, main panel Fig. 2a), despite considerable variation in focal strain performance (Fig. 1a). For example, in sample M3, average focal strain abundance ranged from 9.7 × 10^6^ to 5.3 × 10^8^ CFU/mL (Fig. 1a), but total *E. coli* abundance was relatively stable (ranging from 4.0 × 10^8^ to 7.5 × 10^8^ CFU/mL, main panel Fig. 2). This stability shows that focal strains achieving higher abundance did not increase overall *E. coli* load but displaced resident *E. coli*, indicating a microbiome-specific finite abundance for this species. Taking focal strain abundance as a proportion of the total *E. coli* population in each microcosm revealed that two strains (S3 and S7) consistently reached a comparable or greater abundance than resident *E. coli* (insets Fig. 2). Furthermore, in the respective microbiome samples of these two strains (M3 and M7, where the original untagged strains S3 and S7 are the most abundant resident *E. coli* strain; see Methods), all other focal strains reached relatively low frequencies (insets Fig. 2). This pattern supports the idea that in gut microcosms, S3 and S7 coexisted without strongly affecting each other, while suppressing other *E. coli* strains. The observed strain differences could potentially be explained by variable growth capacities in abiotic conditions, but we found no evidence for this: S3 and S7 showed no advantage in pure culture or pairwise competition in the basal medium used to prepare the microcosms (Supplementary Figs. 4 and 5), nor did they exhibit a particularly broad niche across 95 single-carbon-source conditions (Supplementary Figs. 6 and 7). In contrast, we found some evidence consistent with a role for direct antagonistic interactions between competing *E. coli* strains: S3 and S7 showed inhibitory activity against some other strains on agar (Supplementary Figs. 8 and 9). This phenotype was associated with carriage of a ColE1 plasmid encoding colicin E1 and the corresponding immunity gene. Transfer of the ColE1 plasmid to a susceptible strain (S2) conferred immunity and inhibitory capacity on agar (Supplementary Fig. 10). However, this did not enhance growth performance of S2 in live microbiomes (Supplementary Fig. 11a). This shows genetic elements encoding bacteriocin-mediated antagonism are circulating among resident *E. coli* strains and may contribute to strain-level performance, although the net effects were weak in this manipulation experiment in a distinct genetic background. In summary, strain performance was constrained by a microbiome-specific finite *E. coli* abundance and by intraspecific competition independent of intrinsinc growth capacities.

**Fig. 2:**
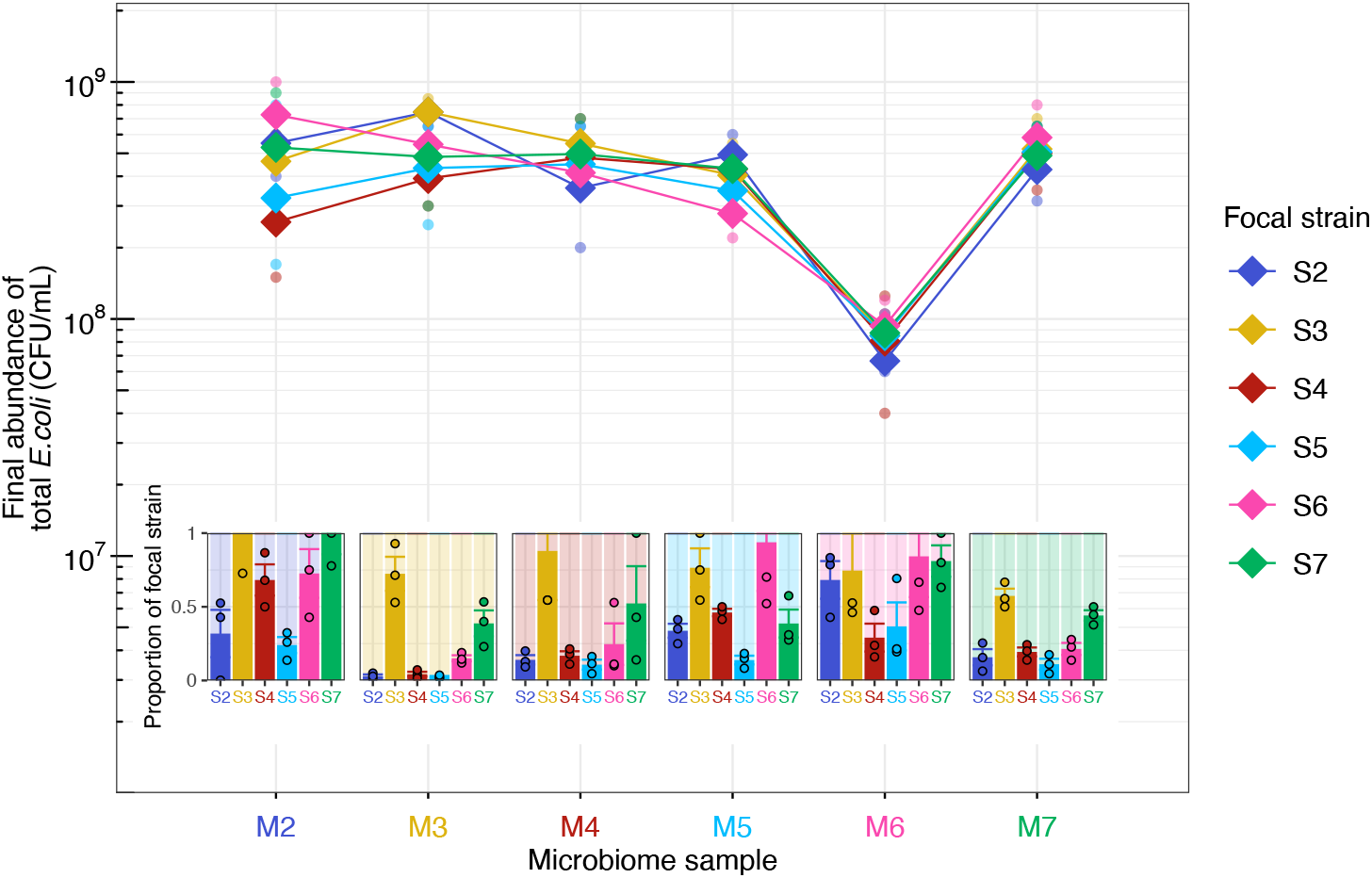
Total *E. coli* abundance and proportion contributed by focal strains in human-gut microcosms. Main panel: final abundance of total *E. coli* (including focal and resident populations) for each live microcosm (Fig. 1a), estimated by plating on chromatic agar without antibiotic. Dots show individual replicates, diamonds represent the average in each combination (geometric mean), and lines connect averages for each focal strain across microbiome samples. Insets: proportion of total *E. coli* represented by the focal strain, obtained by dividing the focal strain’s absolute abundance (Fig. 1a) by the corresponding total *E. coli* abundance (main panel of this figure) for each replicate. Dots show individual replicates and bars/error bars show mean/standard error for each focal strain-microbiome combination. The y-axis scale is limited to 1; individual dots, bars and sections of error bars above 1 due to noise of the method are not displayed. The background colors in lighter shades represent the corresponding most abundant resident strain (e.g., yellow background in microbiome sample 3, M3, corresponds to the original untagged strain 3, S3).

### Interspecific interactions involving acidification contribute to microbiome-specific finite *E. coli* abundance

Aside from resident *E. coli*, differences in the composition or abundance of other resident taxa may alter ecological interactions that determine the success of the incoming strains. Consistent with this possiblilty, we observed substantial variation in taxonomic composition among samples prior to cultivation in microcosms (Fig. 3a, Supplementary Fig. 12). In particular, microbiome sample M6 had a high proportion of Enterococcaceae (27%, brown, Fig. 3a) —not detected in other samples except M4, where its relative abundance was only 0.3%— and a relatively high proportion of Bacteroidaceae compared to other samples (26% vs. mean ± s.d.: 7% ± 3% in other samples, lightest pink, Fig. 3a). M6 also had a relatively low abundance of resident *E. coli* prior to cultivation, shown by both 16S sequencing (all reads assigned to Enterobacteriaceae were of a single amplicon sequence variant, ASV, assigned to *E. coli* and made up 0.02% in M6 vs. mean ± s.d.: 0.06% ± 0.01% in other samples; Supplementary Fig. 12b) and plating (Supplementary Fig. 13a). In terms of total bacterial diversity, M6 was less diverse than other samples (Fig. 3b). The distinctiveness of this sample was further supported by final abundances after our main experiment: total bacterial abundance was approximately 50% lower in microcosms produced with M6 compared with other samples (measured by flow cytometry, Fig. 3c). This trend was even stronger for *E. coli*, in that final total *E. coli* abundance was approximately six-fold lower in microcosms produced with M6 compared with other samples (Fig. 2, Supplementary Fig. 13b), and average focal strain performance was lowest in M6 (Fig. 1a). Thus, microcosms prepared with microbiome sample M6 were the least favourable for microbial growth and particularly for *E. coli*, including our focal strains.

**Fig. 3:**
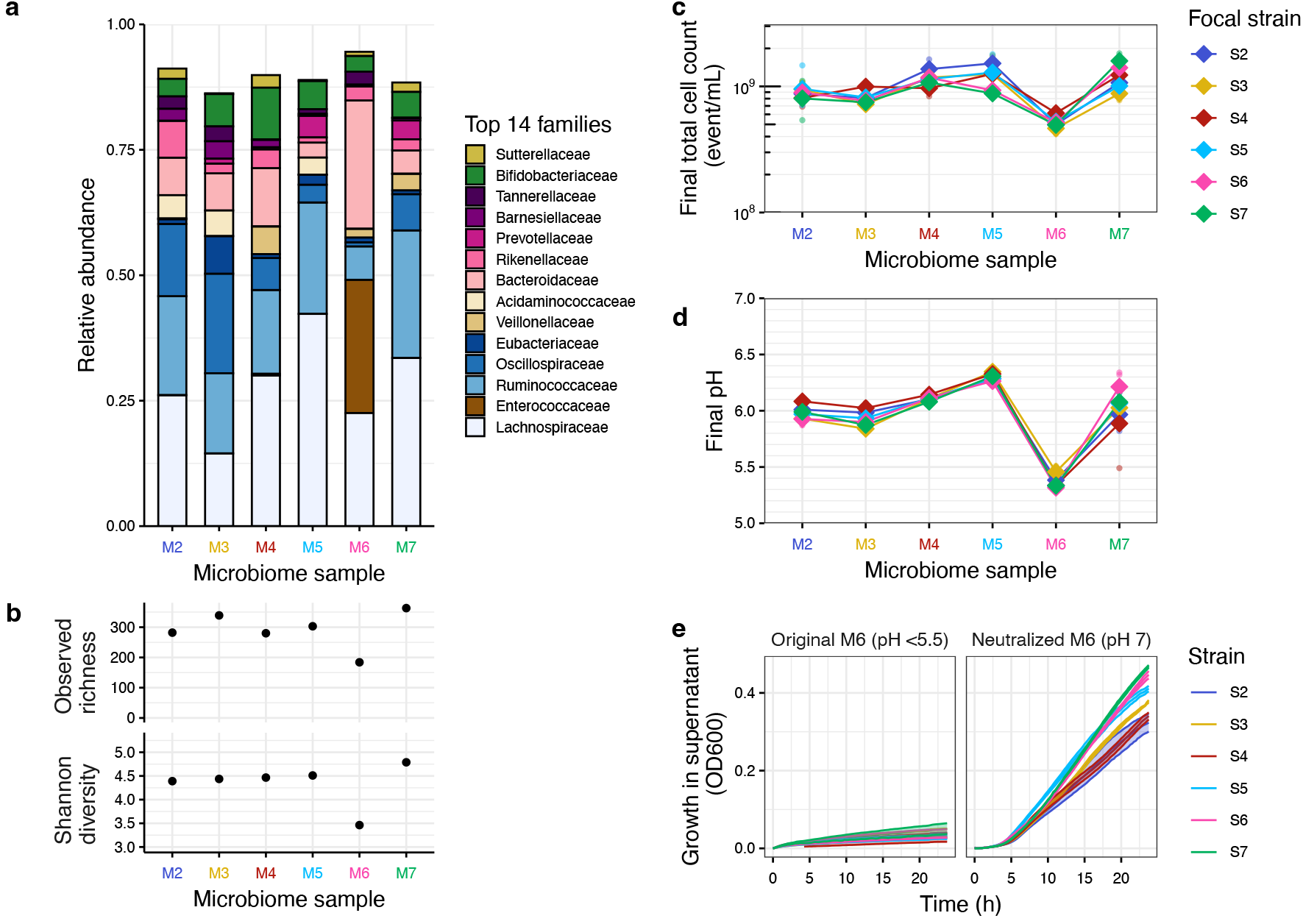
Taxonomic composition and pH-mediated growth constraints. **a** Taxonomic composition of microbiota from the six samples before cultivation. Relative abundance of the fourteen most abundant families across samples, ordered and colored based on phylum (from top to bottom: Proteobacteria in yellow, Actinobacteriota in green; Bacteroidota in pink shades; and Bacillota, further distinguished by class: Negativicutes in beige shades, Bacilli in brown and Clostridia in blue shades). Within phylum, families are ordered by ascending abundance. **b** Alpha diversity (Observed richness and Shannon diversity index) based on ASV-level data for each microbiome sample before cultivation. **c** Final total cell counts in the live microcosms estimated by flow cytometry. Diamonds represent the average (geometric mean) of three replicated microcosms (points for each focal strain and microbiome sample combination) and lines connect the average outcome for microcosms prepared with a given focal strain across microbiome samples. **d** Final pH in the live microcosms. These values show a weak correlation with final abundance of focal strains (Supplementary Fig. 16, R=0.33, p=0.049), largely driven by sample M6. **e** Growth of focal strains under aerobic conditions over 24h (approximated by biomass production, i.e. optical density at 600 nm, OD_600_) in the sterilized supernatant of cultivated microcosms (prepared with the microbiome sample M6 and incubated as in Fig. 1, but without inoculating focal strains) and in the corresponding supernatant adjusted to a neutral pH (7.0). Each curve represents the mean ± standard error of the mean (SE) of three replicates, shown as solid lines with shaded areas of the same color. Equivalent growth measurements in supernant from other microbiome samples are presented in Supplementary Figs. 14 and 15.

We next investigated possible causes of the relatively low focal-strain abundances reached in M6 microcosms (Fig. 1a). This effect was not observed in sterile microcosms prepared with sample M6 (Fig. 1b), suggesting that the relatively unfavorable conditions in M6-microcosms emerged during incubation of the resident microbiota. The most striking feature of the M6 resident microbiota was the high abundance of Enterococcaceae (three ASVs all assigned to *Enterococcus faecium*), which produces lactate during fermentation of carbohydrates [40]. This observation led us to hypothesize that acidification of the microcosm environment could contribute to the unfavourable growth conditions. Supporting this, the final pH of M6 microcosms was considerably lower than that of microcosms prepared with other microbiome samples (on average 5.4 in M6 vs. mean ± s.d. 6.1 ± 0.2 in other samples, Fig. 3d). To test for a causal relationship between low pH and reduced focal-strain growth, we collected supernatant from microcosms prepared with each sample (cultivated in the same conditions as the main experiment but without introducing focal strains) and assessed the growth dynamics of each focal strain in these supernatants (Supplementary Figs. 14 and 15). In contrast to supernatants from other samples, which were less acidic (pH>5.5), the supernatant from M6-microcosms (pH<5.5) did not support any focal-strain growth (Supplementary Fig. 14, Fig. 3e, left panel). However, this growth impairment was reversible: neutralizing the pH of the M6-supernatant with sodium hydroxide restored growth of all focal strains (Fig. 3e, right panel). In summary, the distinctive taxonomic composition of the M6 microbiota led to substantial acidification of the microcosm environment, limiting *E. coli* growth (Fig 2). This process helps to explain the microbiome-to-microbiome variation observed in our main experiment (Fig. 1).

### Transplanting a single taxon across microbiome samples predictably alters resident microbiome-level properties, including *E. coli* growth

We then sought to use the distinctive M6 microbiome as a case study to investigate how resident microbiota contribute to the observed variation of *E. coli* growth among samples from different people (Fig. 1a and 2). Screening taxa from cultivated M6 microcosms, we identified a *C. butyricum* strain that acidified the basal medium in pure culture to a degree comparable with that observed in our earlier experiments with the full M6 microbiome sample (see Methods and Supplementary Fig. 17). This species was not detected in cultivated microcosms from other microbiome samples. *C. butyricum* is a spore-forming anaerobe found in the environment and the gut of humans and animals [41]. It produces butyrate and other short-chain fatty-acids (SCFAs), and is increasingly recognized as probiotic species with documented health benefits in animal models and humans (reviewed in [41]). To test whether *C. butyricum* contributed to the reduction in total *E. coli* abundance in M6, and whether similar effects could be reproduced in different microbiome backgrounds, we transplanted the isolated strain into gut microcosms prepared with the other five samples (M2, M3, M4, M5, M7). Compared with control microcosms (prepared with the same samples but not inoculated with *C. butyricum*), addition of *C. butyricum* markedly reduced final pH of the microcosms (F_1,20_=1477.9, p<0.001), reaching levels comparable to M6 (Fig. 4a). Acidification occured in four of the five tested samples (interaction term: F_4,20_=175.9, p<0.001; significant negative effect for all samples except M4 by post-hoc comparisons using estimated marginal means) and was accompanied by shifts in fermentation product profiles (Fig. 4b). Across all microbiome samples, inoculation with *C. butyricum* led to substantial increases in butyrate concentration relative to control microcosms (+14.8 mM on average, t_4_ = 6.28, p<0.001), and decreases in acetate and lactate concentration (acetate:–8.08 mM on average, t_4_ = –4.37, p = 0.012; lactate: –6.45 mM, t_4_ = –2.33, p = 0.080). Butyrate production in gut microcosms inoculated with *C. butyricum* even exceeded that of *C. butyricum* pure cultures (17.7± 5.4mM vs. 7.0 mM, Supplementary Fig. 18) and among control microcosms, butyrate production was higher in M6 microcosms than microcosms prepared with other microbiome samples (12.9 mM vs. mean ± s.d 2.5 ± 1.8mM, Supplementary Fig. 18). Collectively, these results indicate that *C. butyricum* played a key role in sample M6’s functional profile, and its addition to other microbiome samples was sufficient to establish similar abiotic conditions.

**Fig. 4:**
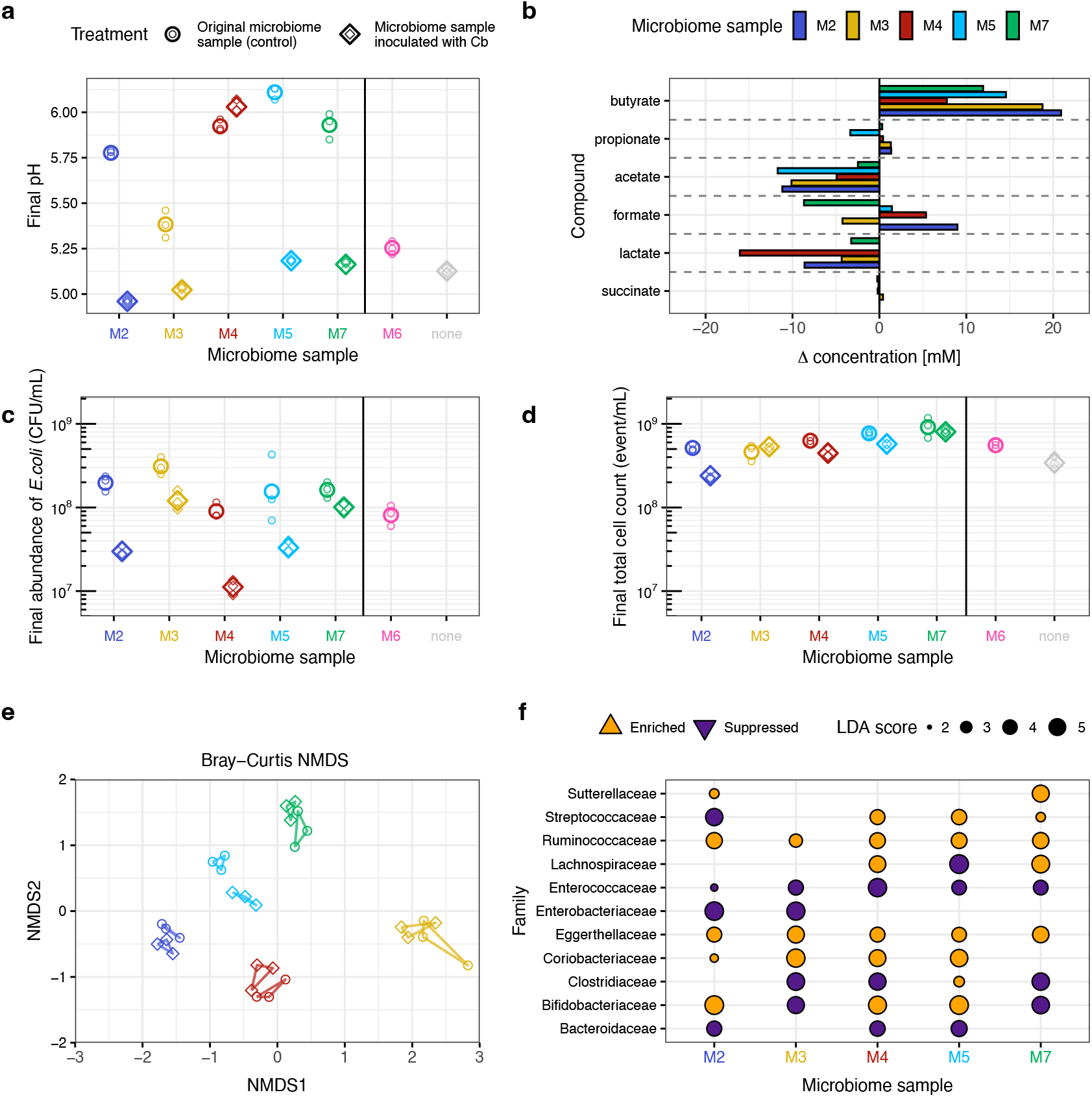
Effects of *C. butyricum* transplantation in human-gut microcosms. **a** Final pH of gut microcosms cultivated with original microbiome samples (without addition of *C. butyricum*; round symbols, left) and the same microbiome samples inoculated with *C. butyricum* (diamond symbols, right). Small points show individual replicates; larger points show geometric means. Two additional controls are shown to the right of the black line: the original M6 microbiome sample, in which *C. butyricum* was naturally present (pink) and microcosms prepared without any microbiome sample (labelled none), but with *C. butyricum* in pure culture (grey). **b** Changes in concentrations of six fermentation products in supernatants of cultivated microcosms with microbiome samples inoculated with *C. butyricum* relative to equivalent microcosms without addition of *C. butyricum* (as in (a)). Bars show differences between the two treatment averages for each compound and microbiome sample. Individual replicates are shown in Supplementary Fig. 18. **c** Final *E. coli* abundance (estimated by plating on chromatic agar) in gut microcosms cultivated with the original microbiome samples uninoculated and the same microbiome samples inoculated with *C. butyricum* (as in (a)). **d** Final total cell counts (estimated by flow cytometry) in gut microcosms cultivated with the original microbiome samples and the same microbiome samples inoculated with *C. butyricum* (as in (a)). **e** Non-metric multidimensional scaling (NMDS) based on Bray-Curtis distances of ASV-level abudances in cultivated gut microcosms. Reads attributed to

We next asked whether addition of *C. butyricum* affected other bacteria. This showed transplantation of *C. butyricum* into other microbiome samples suppressed *E. coli* growth (F_1,20_=117.53, p<0.001, Fig. 4c), to a variable extent depending on the sample (interaction term: F_4,20_=5.45, p=0.0034; significant negative effect in M2, M4, M5 by posthoc tests). Note the most abundant resident *E. coli* strain in each sample corresponded to one of the five focal strains from the main experiment (Fig. 1). Addition of *C. butyricum* also reduced total bacterial growth on average (F_1,20_=25.99, p<0.001, Fig. 4d), again with microbiome-specific effects (interaction term: F_4,20_=7.39, p<0.001; significant negative effect in M2, M4, M5 by posthoc tests). Profiling of community composition with 16S rRNA sequencing revealed that taxonomic composition was also altered by addition of *C. butyricum* (Fig. 4e, PERMANOVA after removing reads attributed to *C. butyricum*, F(1,20) = 37.64, R^2^ = 0.069, p = 0.001), with microbiome-specific effects (interaction term: F(4,20)= 8.04, R^2^ = 0.059, p =0.001). Consistent with this, linear discriminant analysis effect size (LEfSe [42]) revealed multiple taxa that were either enriched or suppressed upon *C. butyricum* supplementation, in terms of relative abundance within each microbial community compared with the corresponding control microcosms and after excluding reads assigned to *C. butyricum* (Fig. 4e, Supplementary Fig. 19). Some taxa were affected consistently across samples; notably the lactate-producing Enterococcaceae family was suppressed, whereas the Ruminococcaceae family, including some well-known butyrate producers such as *Faecalibacterium prausnitzi* [40], was enriched (Fig. 4e, Supplementary Fig. 19). Other taxa exhibited variable responses depending on the microbiome background, such as Lachnospiraceae and Bifidobacteriaceae. Despite a low inoculum chosen to mimic its natural abundance in the original M6 microbiome (see Methods), *C. butyricum* established a substantial population in all inoculated microcosms (28.9% ±11.1% of the community, Supplementary Fig. 20). Overall, these results show *C. butyricum* suppressed growth of some resident bacteria, including *E. coli*, and reshaped community stucture, with some effects consistent across human microbiome samples and others microbiome-specific.

## Discussion

Our reciprocal transplant experiment showed that both absolute and relative population growth of focal *E. coli* strains varied among microbiome samples from different individuals. Intraspecific competition and the resulting displacement of conspecifics shaped strain performance within microbiome samples, each with an inherent finite total *E. coli* abundance. In turn, this variation in total *E. coli* abundance among samples was explained by interspecific interactions involving acidification. We identified a strain of *C. butyricum* contributing to this effect and showed that its transplantation into other microbiome samples reproducibly alters abiotic conditions and biotic interactions.

An important implication of our results is that average growth of commensal *E. coli* varies among samples from different people, with acidification emerging as one driver of this variation. This suggests inter-individual microbiome variation is a key driver of susceptibility to colonisation by incoming *E. coli* and its relative abundance compared to other taxa. This matters because, although individuality in terms of taxonomic composition of human microbiomes is widely documented [8,43], the functional consequences of this are less understood, aside from evidence that dysbiosis can promote colonization by pathogens such as *Clostridioides difficile* [44]. Here, we demonstrate that an even more common form of microbiome variation, among samples from healthy individuals, also has functional consequences. In sample M6, resident microbes acidified the local environment below pH 5.5, largely overcoming the buffering capacity of the basal medium, and inhibiting *E. coli* growth. Microbially driven acidification and its downstream impacts on the growth of other taxa are common in natural microbial communities [45], including in the gut, where some taxa contribute to mild acidification (pH 5.5-6), while others, including *E. coli*, thrive under close-to-neutral conditions (pH 6.5-7) [46,47]. The extent and dynamics of acidification probably differ among individual microbiomes, due to physiological and dietary differences [48], and between spatially and temporally heterogeneous natural settings and our microcosm system. Nevertheless, our findings suggest acidification may be a common mechanism by which person-to-person microbiome variation translates to variable colonization outcomes.

A second implication of our findings is that average *E. coli* growth in a given microbiome can be strongly influenced by a single taxon. Though our study was not designed to explicitly test the link between diversity and colonization resistance, microbiome sample M6 had the lowest diversity among our samples (Fig. 3b), yet showed the stongest suppression of incoming *E. coli* strains, an effect we reproduced by transplanting *C. butyricum* into other microbiome samples. This indicates that functional features, rather than diversity per se, can be decisive in determining colonization outcomes. In turn, this has potentially promising implications in the context of microbiota-oriented health interventions. *C. butyricum* has already emerged as a next-generation probiotic [49,50], with strains such as MIYAIRI 588 used in Asia for decades. However, the ecology of this species remains incompletely characterized. Our results provide new insights in this context in three ways. First, the *C. butyricum* strain we isolated was naturally circulating and identified serendipitously upon microbiome screening, suggesting such interspecies interactions can occur inherently and contribute to inter-microbiome differences even without probiotic intervention. Second, these interactions would remain invisible in sequencing-based surveys. Consistent with this, a recent metagenomic survey of natural Enterobacteriaceae co-excluder taxa [27] did not identify *C. butyricum* among these taxa, despite detecting an overall enrichment in metabolic gene clusters involved in SCFA production. This could reflect both the typically low abundance of this taxon (no ASV assigned to Clostridiaceae in our sample prior to cultivation) and its relatively low prevalence across individuals [51], underscoring how functional consequences of individual-specific taxonomic composition may be missed in sequencing-based analyses. Meanwhile, our microcosm set-up allowed us to detect the functional impact of this species upon cultivation, but it may fail to detect interactions mediated by more fastidious taxa. Third, our replicated and controlled microcosm approach enabled us isolate *C. butyricum’s* impact on other microbes in healthy-human samples, which is not readily disentangled in animal models [52– 54] and human intervention studies [55,56]. This revealed increased butyrate production and suppression of taxa such as Enterobacteriaceae and Enterococcaceae, pointing to a role in colonization resistance [57,58].

A further key finding is that, within microbiome-specific finite *E. coli* abundance, the relative performance of strains varied among samples, while the strain rank order remained generally consistent. This consistency of rank order, resulting in a lack of detectable signature of local adaptation, advances our understanding of the extent to which strains are matched to individual microbiomes. Our results are inconsistent with the idea that the most abundant resident *E. coli* strain from a given microbiome performs relatively well in that microbiome compared to ‘foreign’ strains, or that the home microbiome of a given strain is relatively favourable for that strain compared to other microbiomes. We do not exclude that stronger signatures of local adaptation may be found in other species, such as strict anaerobes with more limited dispersal than aerotolerant microorganisms [59]. Future work expanding tests for local adaptation in human-associated microbiomes may reveal pattern similar to those observed in other ecosystems, where signatures of local adaptation are sometimes detected [23–25] and sometimes not [21,22].

Our results provide new information about the ecological processes driving competition among *E. coli* strains in human-associated microbial communities. Previous work with mice and synthetic communities suggests that incoming strains are suppressed if resident microbes block access to nutrient niches [10,60]. In our system with microbial communities sampled directly from humans and thereby accounting for some interactions and complexity not captured by other approaches, we found no positive association between incoming strain’s ecological success and its metabolic profile relative to the most common resident conspecific (Supplementary Fig. 7). While nutrient competition may play a central role in other experimental conditions with defined limiting nutrients, our results suggest that other mechanisms may have a stronger influence over strain-level *E. coli* success in complex human-associated communities. One such mechanism is direct antagonism mediated by colicin production. Successful strains produced colicin E1, achieving detectable inhibition of some competitors. However, acquiring the plasmid and the corresponding inhibitory phenotype did not boost ecological success of another strain in live microcosms (Supplementary Fig. 10). This does not rule out colicin E1 contributing to the ecological success of strains S3 and S7, but highlights the need to investigate the conditions in which colicins are deployed by *E. coli* populations of different genetic background competing in the gut.

Finally, our results open up new questions about the impact of *E. coli* population structure on colonization success of different gut microbiome samples by individual strains. For example, it is possible that in nature some variability of ecological performance among strains is linked to their phylogenetic relatedness, with more closely related strains potentially sharing some phenotypes [61]. Our results are in some ways consistent with such a trend, in that the two relatively successful strains S3 and S7 belong to the same phylogroup and share some phenotypes (metabolic profile and direct inhibition). However, we would need a much larger number of strains to properly assess the role of phylogenetic structure in colonisation of human samples. Furthermore, although working with samples from different people allowed us to investigate drivers of ecological success among independent human-associated microbial communities, a different experimental design and scale would be needed to quantify the contribution of the identity of the donor to the observed sample-to-sample variation. To do this, more than one sample per individual would be required and, ideally, metadata to allow associations with host factors such as age, diet or immune status to be tested.

In conclusion, our study shows that natural variation in the human gut microbiome has functional consequences, shaping ecological interactions and the success of incoming bacterial strains. While this variation complicates predictions of colonization outcomes, we identified a transferable interspecies interaction that controls *E. coli* growth. These findings improve our understanding of how resident microbes shape colonization outcomes and provide a foundation for microbiome-targeted interventions.

## Material and methods

### Human microbiome samples

Stool samples were collected at the Department of Environmental Systems Science of ETHZ on the 17^th^ of March and the 24^th^ of Mai 2022 from anonymous above-18-year-old healthy donors, who had a body mass index between 19 and 25, were not recovering from surgery, had not taken antibiotics in the previous six months, and did not test positive for SARS-CoV-2 in the previous two months. The sampling protocol was approved by the ETHZ Ethics Commission (EK 2020-N-150). Each volunteer provided one stool sample in a sealed box supplied with an AnaeroGen 2.5L sachet (Thermo Scientific) to maintain an anaerobic atmosphere. We processed the samples within six hours under anaerobic conditions in a vinyl chamber (Coy Labs, USA), containing 2.5 % H_2_, 5% CO_2_ and N_2_ for the rest. We resuspended stool samples 10% *w*:*v* ratio in peptone wash, prepared as previously described (the following components in water, in g per liter: 1.0 peptone from casein, 0.5 L-cysteine, 0.5 bile salts and 0.001 resazurin) [39]. We then froze the resulting fecal microbiome samples with 10% glycerol and stored them at −80°C. For this project, we selected a subset of six fecal microbiome samples, which include a resident microbiota and abiotic components [15] and are referred to as ‘microbiome samples’ hereafter. Because we were unable to isolate *E. coli* from one sample (M1), we excluded it and replaced it with another (M7). Consequently, we performed all experiments with microbiome samples M2-M7. This project developed in parallel in the same laboratory with the project described in Leon-Sampedro et al. [62], which addressed different questions with clinical antibiotic-resistant strains. We used the same sample collection, and microbiome samples M2, M3, M4 in this study correspond to Donor1, 2, 3 in that work.

For 16S rRNA gene sequencing, we sent 2 mL aliquots from each sample to Microsynth AG, Switzerland, for DNA extraction, amplification of the V3V4 region of the 16S rRNA gene using primers 341F (5’-CCTACGGGNGGCWGCAG-3’) and 805R (5’-GACTACHVGGGTATCTAATCC-3’) [63], and sequencing on an Illumina MiSeq instrument. We analyzed the data in R, using the package DADA2 v. 3.18 [64]. Following the workflow described in [65], we filtered and trimmed reads and inferred amplicon sequence variants (ASVs) against the SILVA nr.99 v.138 dataset. After filtering, we obtained a total of 81’503 ± 10’037 (mean ± s.d.) reads assigned to ASVs per sample. We imported the obtained ASV table and corresponding taxa table into phyloseq v 1.46 for downstream analyses and visualizations. We quantified alpha diversity using the function *plot_richness* (measures=*Observed, Shannon*).

### Focal *E. coli* strains

To isolate the focal *E. coli* strains from the samples, we plated fecal microbiome samples at several dilutions on chromatic Mueller-Hinton (MH) agar (Liofilchem, Roseto degli Abruzzi, Italy). From each sample, we chose randomly one purple colony, inoculated it in lysogeny broth (LB, Sigma-Aldrich) and added 25 % glycerol for preservation at −80 °C. We confirmed that picked colonies were *E. coli* by amplifying and sequencing a fragment of *dnaJ*, using DN1-1F (5’-GATYTRCGHTAYAACATGGA-3’) and DN1-2R (5’-TTCACRCCRTYDAAGAARC-3’) as primers [66]. Because multiple *E. coli* strains can co-exist in the same microbiota [7,67], we tested whether the picked colony isolates represented the most abundant *E. coli* strain in each sample by repetitive element palindromic PCR (rep-PCR), using primers REP1R (5’-NNNGCGCCGNCATCAGGC-3’) and REP2R (5’-ACGTCTTATCAGGCCTAC-3’) [68]. This approach allowed us to establish the fingerprint (corresponding to a band pattern on electrophoresis gel) characteristic of each of the six isolated *E. coli* clones and compare it to that of nine other random colonies from each corresponding sample. In five cases, the isolated *E. coli* clone showed a fingerprint matching with 9/9 other colonies from the same sample, and in one case (sample M6) it matched with 8/9 other colonies, indicating that in each sample, the isolated clone represented the most abundant *E. coli* resident strain.

To be able to track these six focal strains (S2-S7) upon inoculation into gut microcosms, where there is a resident microbiota including other *E. coli* cells, we tagged each of them with a selective marker. We did this by electroporating the non-mobilizable plasmid pACYC184 (New England Biolabs), which encodes chloramphenicol resistance [69,70]. Comparing final abundance of each focal strain in sterile microcosms assessed by plating on agar with and without antibiotics (see below) indicated that the plasmid was stable in our experimental conditions (based on pairwise t-tests, abundance differed in only 2/36 combinations, and in different directions: M2, S3 *t*(2) = −8.00, p=0.015 and M3, S7 *t*(2) = 5.64, p=0.030, but both p-values were >0.05 upon adjustment for multiple testing with Bonferroni correction). Furthermore, comparing growth dynamics and competitive fitness of original untagged strains vs. their tagged counterparts in basal medium (Supplementary Fig. 4a and Supplementary Fig. 5b respectively) revealed a small cost of bearing the plasmid, which was comparable among strains and therefore unlikely to confound other causes of strain-to-strain variability in our main microcosm experiments.

We sequenced the whole genome of both the original untagged and the tagged version of the focal strains (Supplementary Methods section **Whole-genome sequencing and analyses**). We also characterized the strains phenotypically, by measuring their metabolic capacity across 96 single-carbon-source environments and testing their inhibitory activity (Supplementary Methods sections **Metabolic profiling across single-carbon-source environments** and **Agar inhibition assays**, respectively). Upon identification of the colicin E1 plasmid as a putative mediator of inhibition, we transferred the plasmid by conjugation to a susceptible strain, to confirm its role in mediating antagonism and test for performance of the resulting transconjugant relative to the original strain in gut microcosms (Supplementary Methods).

### Anaerobic gut microcosms

We quantified growth performance of the focal strains (S2-S7) among microbiome samples (M2-M7) by cultivating them in anaerobic microcosms. We prepared anaerobic 8mL-microcosms in 15mL screw-cap tubes (Sarstedt), containing 0.5 mL microbiome sample (either thawed, or thawed and sterilized by autoclaving for the sterile-microcosms experiment) and 7.5 mL of basal medium, prepared as previously described (resuspending the following components in water, in g per liter: 2.0 peptone from casein; 2.0 tryptone, 2.0 yeast extract, 0.1 NaCl, 0.04 K_2_HPO_4_, 0.04 KH_2_PO_4_, 0.01 MgSO_4_x7H_2_O, 0.01 CaCl_2_x6H_2_O, 2 NaHCO_3_, 1.8 Tween 80, 0.005 hemin, 0.5 L-cysteine, 0.5 bile salts, 2 starch, 1.5 casein, 0.001 resazurin; adjusting the pH to 7; autoclaving and finally adding 0.001 g per liter menadion) [39]. We made three replicate microcosms in each combination of focal strain and microbiome sample, giving 108 microcosms in total for the live- and sterile-microcosm experiments. We added focal strains (tagged versions) after diluting independent overnight cultures (incubated at 37 °C statically and anaerobically in 200 µl basal medium), aiming to achieve an inoculum density between 10^4^ and 10^5^ CFU/mL. We chose this inoculum density to start with comparable amounts of focal and resident *E. coli* (Supplementary Fig. 13a). We incubated microcosms at 37 °C, statically and anaerobically for 24 h. This single 24-h growth cycle was selected based on our previous findings [39], which showed that this timeframe captures variation in focal strain population growth while maintaining the dominant family-level structure of the original inoculum. Although fast-growing facultative anaerobes (particularly Enterobacteriaceae and Enterococcaceae) are predictably enriched under these conditions [71], the short cultivation period preserves ecologically relevant community interactions.

We assessed abundance of focal strain and total *E. coli* by sampling 200 µl from well-mixed microcosms and spot-plating dilutions on chromatic MH agar plates supplemented with 25 µg/mL chloramphenicol and without antibiotics respectively. For all microcosm experiments, we supplemented aliquots or the whole volume of the cultivated microcosms with 10% glycerol and stored them at −80 °C for further use.

In live microcosms, effective starting densities varied among focal strains (F_5,72_=52.85, *p*<0.001, ranging between 2 × 10^4^ and 8 × 10^5^ CFU/mL, Supplementary Fig. 21a). A separate experiment, in which we inoculated one focal strain at different densities into three different microbiome samples, indicated that within this range, final abundance was independent of inoculum density (R=0.23, p=0.25, Supplementary Fig. 21b). Therefore, we took final abundance as a measure of focal strain growth performance. For statistical analysis, we applied analysis of variance (ANOVA, aov in R) with strain, microbiome, and their interaction as factors, and log-transformed CFU counts as the response variable. For both live- and sterile-microcosm experiments, we quantified average local adaptation, defined as the sympatric-allopatric contrast [17] (Supplementary Methods section **Quantifying local adaptation**).

We estimated total bacterial abundance in aliquots from live microcosms from the main experiment using flow-cytometry. We thawed aliquots of the 108 microcosms, diluted them 1:1000 in 200 µl phosphate-buffered-saline (PBS) and stained cells with the addition of SYBR Green (Invitrogen, Thermo F. Scientific). We used a Novocyte 2000R (ACEA Biosciences, San Diego, CA, USA) with the following parameters: 50 µl flow rate, 14.6 core diameter and 100’000 events recorded per sample. We measured pH in supernatant from the 108 microcosms from the main experiment, after thawing and centrifuging them, using a portable pH-meter FiveGo F2 (Mettler Toledo).

In a separate experiment, we estimated initial and final abundance of resident *E. coli* in the absence of tagged focal strains (Supplementary Fig. 13), preparing and cultivating microcosms in the same way as we did for the live microcosm experiment, but without inoculating focal strains. Here as well, we made three parallel replicates, giving a total of 18 microcosms. We made pH measurements by thawing the 18 cultivated microcosms and using the pH-Fix 2.0-9.0 test-strips (Macherey-Nagel). From the same microcosms we prepared supernatant, removing bacterial cells by centrifugation (4000 g, 10 min) and sterile filtering with a pore size 0.2 µm (Filtropur S, Starstedt). We adjusted the pH of each supernatant by adding few microliters of 1M sodium hydroxide and remeasuring, until reaching a pH of 6.5-7 (up to 6 mM sodium hydroxide in the three supernatants from M6 microcosms).

### Growth dynamics of focal strains

We monitored population growth dynamics of each focal strain in various conditions (basal medium, sterile microcosm conditions and supernatant of cultivated microcosms) and always aerobically. To do so, we diluted independent overnight cultures (incubated at 37 °C shaking and aerobically in 200 µl basal medium) 1:1’000 in 200 µl of the corresponding media in a microplate. We then incubated microplates at 37 °C in a plate reader (Tecan NanoQuant Infinite M200 Pro), shaking and measuring optical density at 600 nm (OD600) every 15 min and during 24 h. To assess contribution of strain identity and selective marker (pACYC184) to the variability of growth dynamics, we monitored growth of both tagged and untagged version of each focal strain in basal medium (Supplementary Fig. 4a). In all other conditions, we characterized growth of the tagged version of focal strains exclusively. To monitor growth in sterile microcosm conditions (Supplementary Fig. 4b and Supplementary Fig. 14), we prepared one tube per sample containing 0.5 mL of autoclaved microbiome sample and 7.5 mL of basal medium, and we let particles sediment overnight at 4 °C. We then inoculated each focal strain in each sterile microcosm medium, with three replicates for each strain-microbiome combination. To monitor growth across supernatants of cultivated microcosms (Fig. 3e, Supplementary Fig. 14), we inoculated each focal strain into three replicated supernatants from each microbiome sample, prepared as described above.

To estimate focal strains’ fitness in different conditions, we estimated maximal growth rates and area under the curve from all pure culture growth curves and to quantify relative competitive fitness we performed a pairwise competition experiment in basal medium (Supplementary Methods section **Quantifying absolute and relative competitive fitness of focal strains**).

### *C. butyricum* isolation and supplementation experiment

We plated several dilutions of cultivated microcosms on TPY broth agar (Condolab) under anaerobic conditions and incubated them for 40h. We inspected colony morphology on plates from M6 microcosms and picked all colonies with a visibly distinct morphology. We submitted those isolates to MALDI-TOF analysis and detected four distinct species: *E. coli, E. faecium, Bifidobacterium longum* and *C. butyricum*. We subsequently cultured each isolate in basal medium and measured pH of the cultures at an endpoint with pH-meter FiveGo F2. We found that only the *C. butyricum* acidified basal medium (Supplementary Fig. 17). We stored some sterile supernatant of all replicates of these cultures at 4 °C. We inspected plates from other samples looking for colonies with the characteristic morphology of *C. butyricum* (slightly irregular shape, creamy and notably larger than *E. coli* colonies) but did not detect any. We also revisited the 16S rRNA data from the microbiome samples to query ASVs assigned to the Clostridiaceae family. None were detected in sample M6 and only a small number of reads were assigned to an unknown species within the *Clostridium sensu stricto 1* genus in some other samples (27 reads in M2, 48 in M3, and 10 in M5). We concluded that in sample M6, *C. butyricum* accounted for fewer than 10^4^ CFU/mL (as it was represented by less than one read per 100’000, within a total bacterial load of approximatively 10^9^ CFU/mL).

We performed a supplementation experiment in gut microcosms by preparing triplicate 8-mL anaerobic cultures in basal medium from five microbiome samples (M2, M3, M4, M5 and M7) as described above. For each sample, cultures were either supplemented with *C. butyricum* or left untreated as controls. In addition, we included triplicates of the original microbiome sample M6 as a reference, giving 33 microcosms in total, as well as pure *C. butyricum* cultures. Supplementation was performed by inoculating *C. butyricum* independent overnight cultures diluted 1:1’000. We incubated microcosms for 24 h and estimated *E. coli* abundance by plating on chromatic agar and total bacterial abundance with flow cytometry as described above. We also measured pH and stored sterile filtered supernatants at 4 °C for further characterization. For statistical analysis, we applied analysis of variance (ANOVA, aov in R) with treatment, microbiome, and their interaction as factors, and log-transformed CFU counts as the response variable. We used the emmeans package to compute estimated marginal means and performed pairwise comparisons of treatments within each microbiome.

### Fermentation products and taxonomic profiling of cultivated microcosms

We quantified six key fermentation products (succinate, lactate, formate, acetate, propionate, and butyrate) in culture supernatants of the 33 cultivated microcosms and pure cultures using high-performance liquid chromatography (HPLC). We injected 20 µL of each sample into a Phenomenex Rezex ROA–Organic Acid H? (8%) column at 40 °C, using 2.5 mM H2SO_4_ in water as the mobile phase with a flow rate of 0.4 mL min^−1^, as described in [72]. To prepare calibration curves and define the expected retention times, we injected metabolite mixtures with each metabolite at concentrations of 1, 5, 10, 15, and 20 mM (sodium salts, Sigma-Aldrich). For each standard and sample, the refractive index detector recorded signals over 40 min, and we exported chromatographic data as detector response (µRIU) versus time (s). We processed the data in Python using SciPy. Implementing the workflow outlined in hplc-py [73], we applied baseline correction (*minimum_filter1d*), detected peaks (*find_peaks*), and fitted multi-Gaussian models (*curve_fit*) accounting for both expected and unexpected peaks. We verified peak fitting accuracy by generating diagnostic plots with Matplotlib. Finally, we integrated peak areas and converted the areas into concentrations using the calibration curves.

We profiled taxonomic composition of the 33 cultivated microcosms from the *C. butyricum* supplementation experiment with 16S rRNA sequencing using the same workflow as for the fecal microbiome samples (see above). Quality control and filtering resulted in 107’836 ± 9339 (mean ± s.d.) reads assigned to ASVs per sample. We used the phyloseq framework in R to aggregate ASVs to family level and summarize community composition with relative abundance barplots (Supplementary Fig. 20). For downstream analyses, we excluded sample M6 and removed reads attributed to *C. butyricum*. We assessed beta diversity using the ASV abundance matrix and performed ordination via non-metric multidimensional scaling (NMDS) with *metaMDS* from the vegan package (v. 2.7-1), specifying Bray–Curtis dissimilarities (distance=*bray*). We visualized differences in microbial communities across treatments and microbiomes with NMDS plots. We tested the effects of treatment and microbiome on community composition using PERMANOVA via the *adonis2* function applied to Bray–Curtis distances. We assessed differential abundance of taxa across treatments using *lefser* from the microbiomeMarker package (v.1.8.0). We performed analyses at different taxonomic levels (Family and Genus) within each microbiome sample separately, identifying taxa with significant LDA scores as enriched or suppressed in response to treatment. Finally, we combined results from all samples for visualization.

## Data availability

Sequences of the genomes of the six *E. coli* strains and 16S rRNA amplicons from the six fecal microbiome samples and from the 33 cultivated microcosms are uploaded to ENA (European Nucleotide Archive) and accessible through project PRJEB80311. The code and all other data supporting the findings of this study will be made available on Figshare (https://doi.org/10.6084/m9.figshare.27854763) upon publication.

## Acknowledgements

We thank the Pathogen Ecology group, Carolin Wendling and Angus Buckling for their feedback. We thank the team at the Clinic for Infectious Diseases and Hospital Epidemiology of the University Hospital Zurich (USZ) for providing access to MALDI-TOF and for their support.

