## Supplementary Information for "Interspecies interaction controls *Escherichia coli* growth in human gut microbiome samples"

### Supplementary Methods

**Focal strains' whole-genome sequencing and analyses**

We sequenced the whole genome of both the original untagged and the tagged version of the focal strains (with short-read sequencing and both short- and long-read-sequencing, respectively). DNA extraction and sequencing were performed at the Institute of Medical Microbiology of the University of Zurich (IMM, UZH). DNA was extracted with the Maxwell RSC Blood DNA Kit (Promega, Madison USA). Libraries were prepared with the QIAseq FX DNA Library Kit (QIAGEN, Hilden, Germany) for 150-paired-end sequencing on an Illumina NextSeq1000, and with the Rapid Barcoding Kit 24 V13 (Oxford Nanopore Technologies, Oxford, UK) for sequencing with the Minion R.9.3 flowcell on a GridION (Oxford Nanopore Technologies, Oxford, UK).

We filtered long reads with Filtlong v 0.2.1 (https://github.com/rrwick/Filtlong) by discarding reads shorter than 1 kb and 10% read bases with poorest quality. We generated hybrid assemblies combining short- and filtered long-reads with Unicycler [1]. Despite having scaffolded the assemblies with long reads, some remained incomplete, i.e. featured uncircularized contiguous sequences (contigs). We extracted assembly statistics with assembly-stats v. 1.0.1 (https://github.com/sanger-pathogens/assembly-stats) (Supplementary File 1). We determined the putative chromosomal or plasmid origin of each contig using PlasmidHunter [2]. We annotated the assemblies with Prokka v 1.14-6 [3] and constructed a pangenome with Roary v 3.13.0 [4], using the command -e mafft. We used the resulting core gene alignment to construct a maximum likelihood tree with FastTree 2 [5]. We determined phylogroup and sequence type of the strains with the tools ClermonTyping (http://clermontyping.iame-research.center) and mlst (https://github.com/tseemann/mlst) (Supplementary Fig. 1).

We screened assemblies for characterized *E. coli* virulence factors (including bacteriocins) with Abricate v 0.5 (https://github.com/tseemann/abricate), using the database *ecoli_vf* (https://github.com/phac-nml/ecoli_vf) and default parameters (–minid 80%). Then, we mined the assemblies for regions encoding putative prophages and bacteriocins clusters with PHASTER [6] and BAGEL4 [7] respectively. To create a list of putative bacteriocins, we considered the areas of interest identified by BAGEL4 that included a core peptide and inspected genomic context and whether core peptides matched functionally characterized *E. coli* bacteriocins. Most of them did match bacteriocin genes included in the *ecoli_vf* database or, in one case, the recently characterized gene cluster mccPDI [8,9]. One core peptide—identified on the chromosome of the strains from phylogroup B2 (S2, S4 and S5)— encoded the T6SS effector protein Hcp fused with extended-toxin domains (Hcp-ET3-4), also found on the chromosome of the reference strain *E. coli* Nissle [10,11]. We excluded this hit from the list of putative bacteriocin because it is not formally classified as such. We used clinker [12] to visualize the alignments of the colicin E1 plasmids carried by S3 and S7 (Supplementary Fig. 8d).

**Metabolic profiling across single-carbon-source environments**

We determined the metabolic profile of each focal strain (original untagged version) by measuring their ability to use 96 carbon sources supplied individually in the wells of Biolog AN microplates (Endotell AG, Switzerland). Under anaerobic conditions, we resuspended the carbon sources of the Biolog AN microplate in 100 µl of the manufacturer inoculation fluid (AN-IF) and inoculated focal strains (original untagged version) by sampling 1 µl from independent overnight cultures (incubated at 37 °C statically and anaerobically in 200 µl LB in microplates) with a 96-pin microplate replicator. We sealed the microplates in an air-tight bag, which we incubated at 37 °C, shaking. After 24 h, we measured OD600 in each well. We profiled metabolism of the six strains in three experimental blocks and for each plate subtracted the value of the blank well from the value obtained in the other wells. We considered a given substrate usable by a given strain if standard error of mean subtracted from the mean was higher than the arbitrary threshold of 0.1 (Supplementary Fig. 6). We defined focal-strain nutrient niche size as the number of usable substrates. We quantified average growth by each focal strain on its niche by computing the mean OD600 across usable substrates. Finally, we assessed focal-strain private niche size in each microbiome sample based on the binarized version of the six metabolic profiles shown on Supplementary Fig. 7a: we computed the number of usable substrates that a given focal strain could use that another strain (representing the most abundant resident *E. coli* in each microbiome sample) could not use.

**Agar inhibition assays**

We tested the inhibitory activity of the focal strains using agar inhibition assays. We prepared bacterial lawns by diluting 200 µl of an exponential-phase culture from the indicator strain (approximately $2 \times 10^8$ colony-forming-units (CFU)/mL) in 6 mL soft LB agar (0.4% *w*:*v* agarose), pouring it onto LB agar plates and allowing it to dry for 30 min. We then spotted 5 µl of filter-sterilized supernatant from stationary cultures of the strains to be tested. We also used a variation of this protocol, using filter-sterilized supernatant of exponential-phase cultures with and without induction of the SOS response, prepared by diluting stationary cultures 1:100 in LB, and incubating for 5h with and without addition of 0.5 µg/mL mitomycin C after 1h [13]. In the reciprocal inhibition assay, we used the focal strains (original untagged versions) as both indicator and producer (Supplementary Fig. 9a). In a separate experiment, to rule out that the observed clearance zones resulted from lysis by phages, we used strain K12-MG1655 lambda+ Δbor::cm [14] as producer in parallel with focal strains S3 and S7 and spotted dilutions of the filter-sterilized supernatants of exponential-phase cultures exposed to mitomycin C (Supplementary Fig. 9b).

In strain S3, the colicin plasmid hypothesized to be responsible for the observed inhibitory phenotype encoded the mobilization protein MbeC and the relaxase MbeA (Supplementary Fig. 8d) and was therefore putatively mobilizable. S3 also carried an IncF conjugative plasmid, which encoded several resistance genes. We performed a conjugation experiment with the tagged version of S2 as recipient and the original untagged version of S3 as donor. We used the natural resistance to trimethoprim conferred by the IncF plasmid and the chloramphenicol resistance of the recipient to select for transconjugants. We selected ten transconjugant colonies and performed colony PCRs to verify that they carried the IncF plasmid (with primers IncFII_F 5′-CCGCATAGAAGCTGTTGCT-3′ and IncFII_R 5′-TCCTGCACTTATGTTGCACAG-3′) and the two original S2 plasmids (IncB/O/K/Z and the IncX3, with primers IncO_F 5′-GTCCGGAAAGCCAGAAAACG-3′ and IncO_R 5′-CCGCCAAGTTCGACAGGAAG-3′ ; IncX3_F 5′-GGGGTAACTCTTGCATCCCTT-3′ and IncX3_R 5′-TTTCAGAGCTGCATAAGAGGCA-3′ respectively). As expected, all transconjugants carried the three plasmids. Furthermore, we screened for colE1 plasmid carriage (with primers colE1_F 5′-AAGCCCGTAAAGAAGCGGAA-3′ and 5′-ATTTCGTTTTCCAGCGAGCG-3′) and found that 1/10 transconjugants also carried the colE1 plasmid. We named this transconjugant S2t1 and, as control to discriminate possible phenotypic effects conferred by acquisition of the IncF plasmid rather than the colE1 plasmid, we picked one of the other transconjugants without colE1 plasmid (S2t2).

**Quantifying local adaptation**

For both live- and sterile-microcosm experiments, we quantified average local adaptation, defined as the sympatric-allopatric (SA) contrast [17] (Supplementary Fig. 2). The SA-contrast corresponds to the difference between average focal-strain performance in sympatric strain-microbiome combinations (the six combinations in which focal strains were inoculated into their home microbiome sample) vs. allopatric strain-microbiome combinations (the other 30 combinations). For each data set, we tested whether the observed SA-contrast value differed from the null expectation (no average difference between sympatric and allopatric combinations), by first calculating the SA-contrast score in each of 1000 random permutations of the same dataset (where the same 36 values as used in calculating the observed SA-contrast were randomly reassigned across strain-microbiome combinations). We then assessed the probability that the observed value could have arisen by chance from the proportion of randomly permuted SA-contrast scores lower than the observed result.

**Quantifying absolute and relative competitive fitness of focal strains**

From all growth curves in pure cultures, we estimated maximal growth rates with the R package growthrates [66] by fitting a linear regression to the 8-datapoint subset (corresponding to a 2-hours window) with the steepest log-linear increase. We extracted the area under the curve, which correspond to a sum of the all 96 OD600 values over 24 h, with the R package growthcurver [66].

To quantify relative competitive fitness of the *E. coli* strains, we prepared triplicates of all strain-strain combinations by diluting independent overnight cultures (incubated at 37 °C shaking and aerobically in 200 µl basal medium) 1:1'000 and adding them to 200 µl of basal medium in a microplate. Thus, the competitors were initially at a 1:1 ratio and an abundance of approximatively $10^5$ CFU/mL and we co-cultivated them during 24 h. Each combination included an untagged and a tagged version of the strains, to enable monitoring of initial and final abundance of focal strains (by enumerating CFU on agar with antibiotics) and estimation of initial and final abundace of untagged strains (by subtracting CFU attributed to focal strain from total CFU enumerated on agar without antibiotics). We then computed the selection rate constant with the formula $ln\,(\frac{A_f}{A_i}) - ln(\frac{B_f}{B_i})$, where A corresponds to the tagged strain population and B the untagged strain population, and the subscripts i and f to initial and final abundance [67].

#### Supplementary Figures

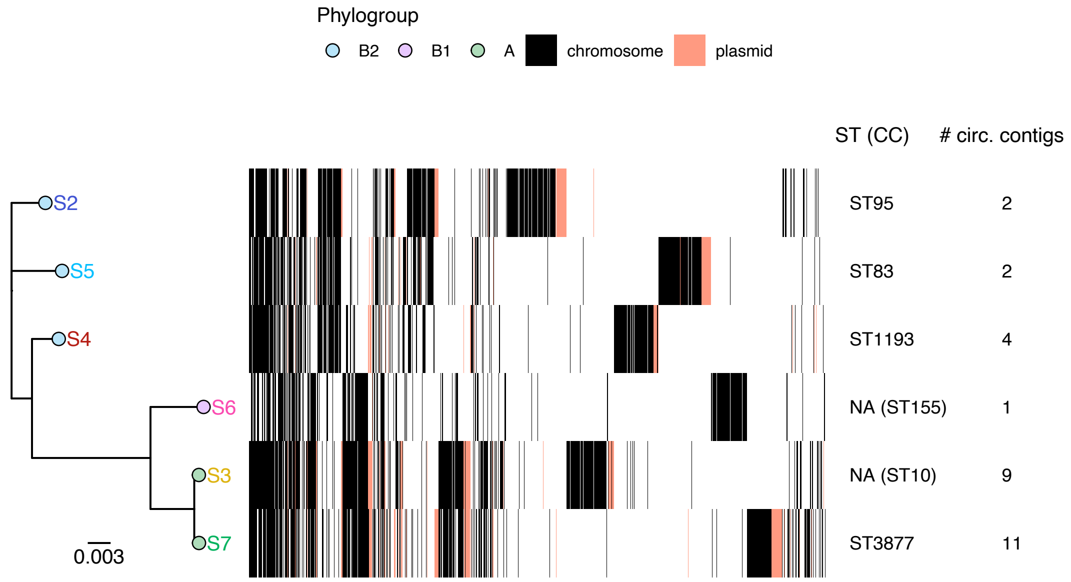


**Supplementary Fig. 1 Phylogeny and accessory genes' map of the six *E. coli* clones isolated from stool samples.** On the left, midpoint-rooted maximum-likelihood tree based on core gene alignment (of length 3.2 Mb, so scale-bar corresponds to ~10'000 substitutions). Color of the tips and their label correspond to phylogroup and color identifying the six *E. coli* isolates throughout the manuscript respectively. On the accessory genes map, every column corresponds to one of the 5'254 accessory genes (i.e., genes not shared among all isolates). A gene absent from a given genome assembly is displayed as white bar on the corresponding row. A gene that is present is displayed as black or coral bar, if putatively located on the chromosome or on a plasmid respectively (see Supplementary Methods). On the right, on the row corresponding to each isolate shown on the tree, sequence types (if not assigned, NA, clonal complex, CC, in brackets) and number of circularized contigs besides the chromosome are displayed. The latter reflect approximatively the number of plasmids, possibly underestimated, because among the contigs left un-circularized in assemblies which weren't completely closed, some were categorized as plasmid content (see Supplementary Methods).

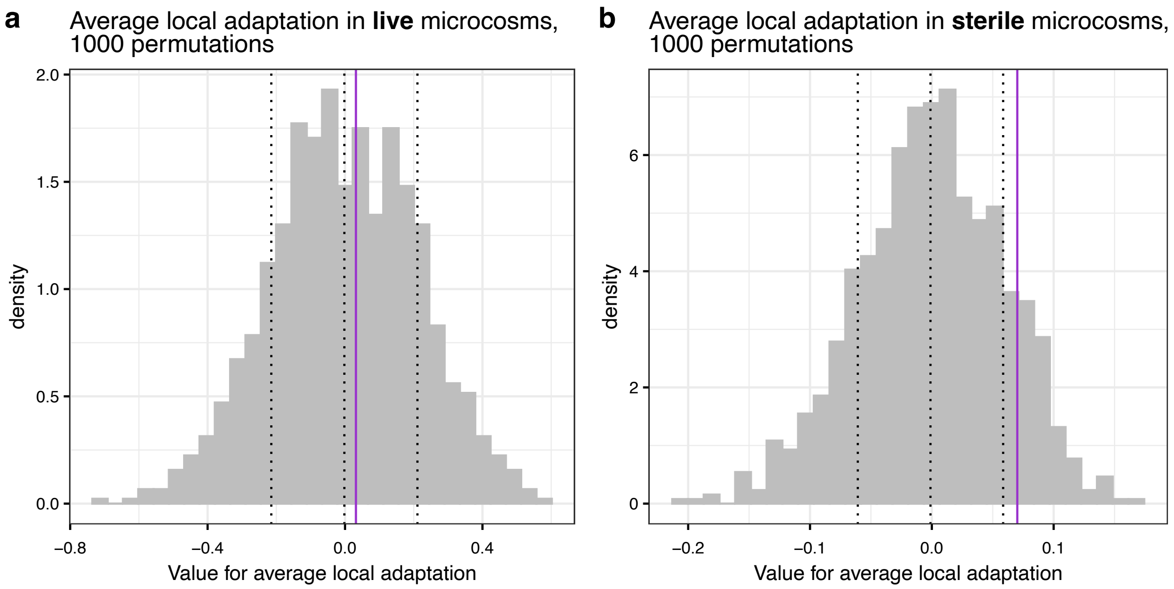


**Supplementary Fig. 2 Quantification of average local adaptation in live (a) and sterile (b) microcosms**. In each panel the observed sympatric-allopatric (SA) contrast value is shown with a purple vertical line overlaid on the distribution of SA-contrast values obtained for 1’000 random permutations of the same dataset (grey histogram). **a** In live microcosms, the observed SA-contrast value is 0.032, and the distribution of values obtained for 1000 permutations has mean ± s.d. 0 ± 0.21. The probability that the observed value could have arisen by chance is 90% , because the shuffled results exceed the true result in 897/1’001 cases. **b** In sterile microcosms, the observed SA-contrast value is 0.070, and the distribution of values obtained for 1000 permutations has mean ± s.d. 0 ± 0.06. The probability that the observed value could have arisen by chance is 25%, because the shuffled results exceed the true result in 251/1001 cases.

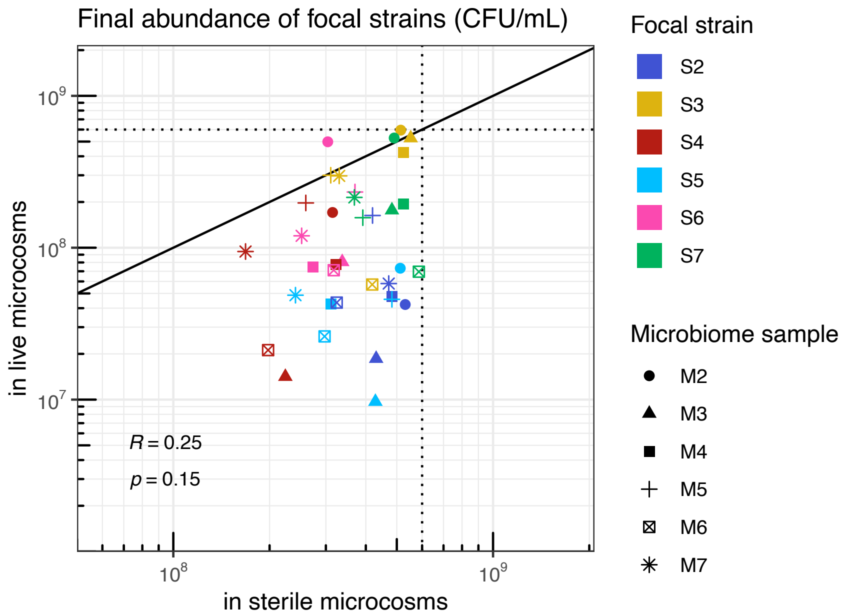


**Supplementary Fig. 3 Correlation between growth performance of focal *E. coli* strains in live vs. sterile microcosms** (represented in Fig. 1a and Fig. 1b respectively). The solid line is the 1:1 slope and dashed lines show the maximal final abundance reached in both experiments (5.9 x $10^8$ CFU/mL). Different colors distinguish the six strains, and different shapes distinguish the six microbiomes. Pearson's correlation coefficient (R) and the corresponding p-value are displayed on the bottom-left corner.

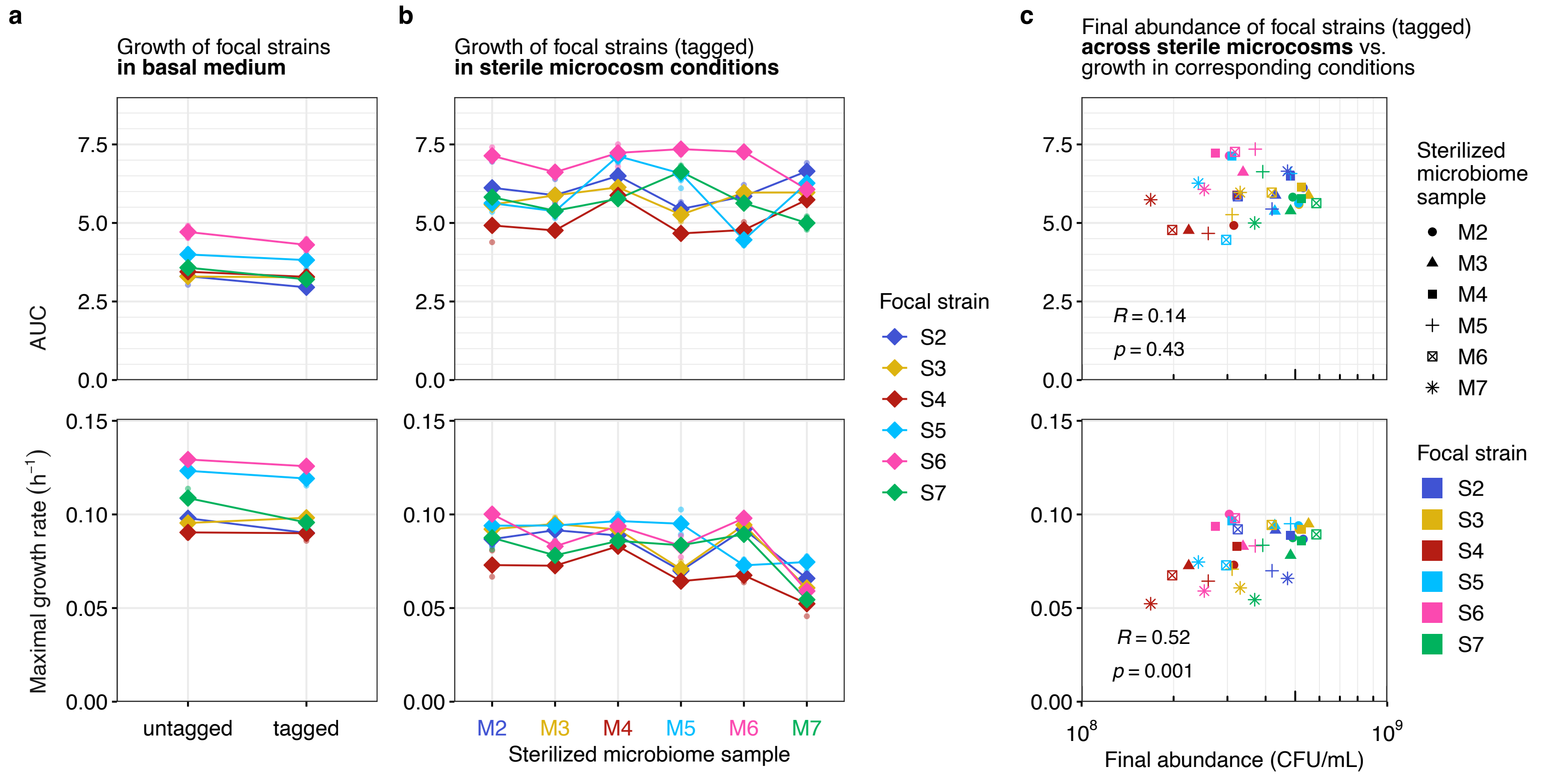


**Supplementary Fig. 4 Growth dynamics of focal strains in basal medium and across sterile microcosm conditions. a** Area under the curve (AUC) and maximal growth rate of original (untagged) and tagged version of each *E. coli* strain (x-axis) during 24h growth in basal medium (in a microplate under aerobic conditions with shaking). Diamonds represent the average of three replicates and dots show individual replicates. To determine whether untagged and tagged version of each strain had different growth dynamics, we carried out pairwise t-tests. Growth dynamics differed only for strain S7, whose tagged version grew slightly worse than the untagged counterpart (AUC: $t(3.72)=3.90$, p=0.020 and maximal growth rate: $t(3.83)=4.05$, p=0.017; but both p-values were >0.05 upon adjustment for multiple testing with Bonferroni correction). **b** Area under the curve and maximal growth rate of focal strains (tagged) during 24h growth in sterile microcosms conditions (i.e. 6.25% *v*:*v* sterilized microbiome sample in basal medium, so equivalent abiotic conditions as experiment from Fig. 1b, but in a microplate under aerobic conditions with shaking). The raw data underlying these plots are shown in bottom panel of Supplementary Fig. 14. **c** Correlation between final abundance of focal strains across sterile microcosms (x-axis, data from Fig. 1b) and their growth dynamics in corresponding abiotic conditions (y-axis, data from panel b of this figure). Different colors distinguish the six strains, and different shapes distinguish the six sterilized microbiome samples. Pearson's correlation coefficient (R) and the corresponding p-value are displayed on the bottom-left corner.

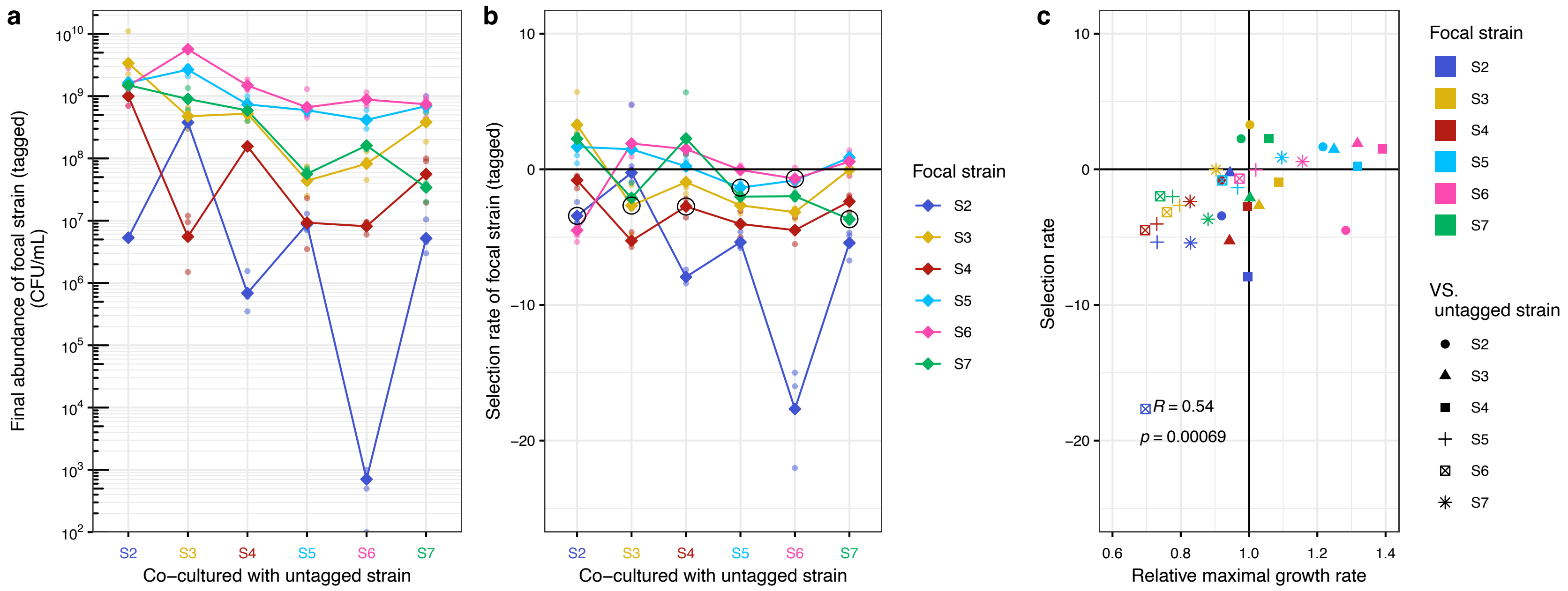


**Supplementary Fig. 5 Reciprocal pairwise competition of focal strains in basal medium.** Tagged focal strains were co-cultured with untagged versions of the same strains in basal medium in a microplate under aerobic conditions with shaking. Final abundance **(a)** and selection rate constant **(b)** of each focal strain (tagged) when co-cultured with the untagged version of each strain, representing the most abundant resident *E. coli* strain in each sample. Diamonds represent the average (geometric mean) of three replicates for each combination and dots show individual replicates. Lines connect the average outcome for each tagged strain across untagged strains. In **b**, selection rate constants above 0 (see reference line), reflect an advantage of the focal strain population against untagged strain population. In combinations where a given focal strain was co-cultured with its untagged counterpart, highlighted with a black circle, the selection rate was slightly below 0 (ranging from -3.7 to -0.7), meaning that for each focal strain, the population of the untagged version had a slight advantage relative to the population of the tagged counterpart. **c** Correlation between relative maximal growth rate (x-axis, calculated from data shown in bottom subpanel of Supplementary Fig. 4a; by dividing the maximal growth rate of the tagged version by that of the untagged version of each focal strain) and selection rate (y-aixs, data from panel b of this figure). Relative growth rates above 1 (on the right of the vertical reference line) reflect a higher maximal growth rate for the focal strain (tagged) relative to the corresponding untagged strain. Pearson's correlation coefficient (R) and the corresponding p-value are shown on the bottom-left corner.

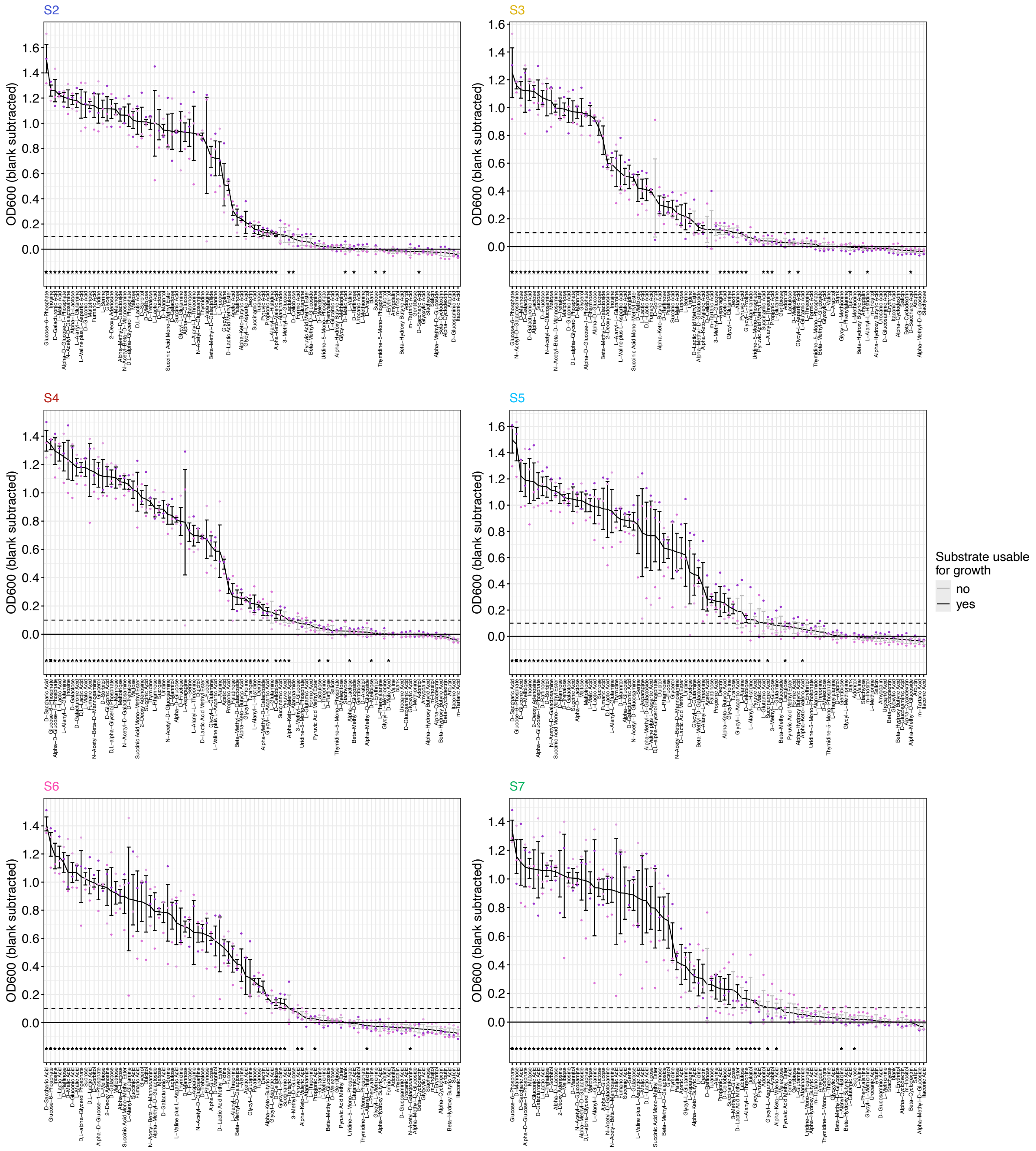


**Supplementary Fig. 6 Metabolic profile of the focal strains.** Raw data underlying Supplementary Fig. 7a. OD600nm reached by each strain (displayed in different panels) across the 95 single-carbon-source environments tested (rearranged within each panel based on mean by descending order). Dots show individual replicates, and the line connects the mean across growth substrates. Error bars represent standard error of mean (SE), and their color highlights the outcome of mean-SE: black if it is above the threshold 0.1 (dotted line) and grey if it is below, with corresponding substrates considered usable by a given strain or not, respectively. Growth substrates usable by at least one of the strains (and thus included in downstream analyses and displayed on Supplementary Fig. 7a) are labelled with a star on the bottom.

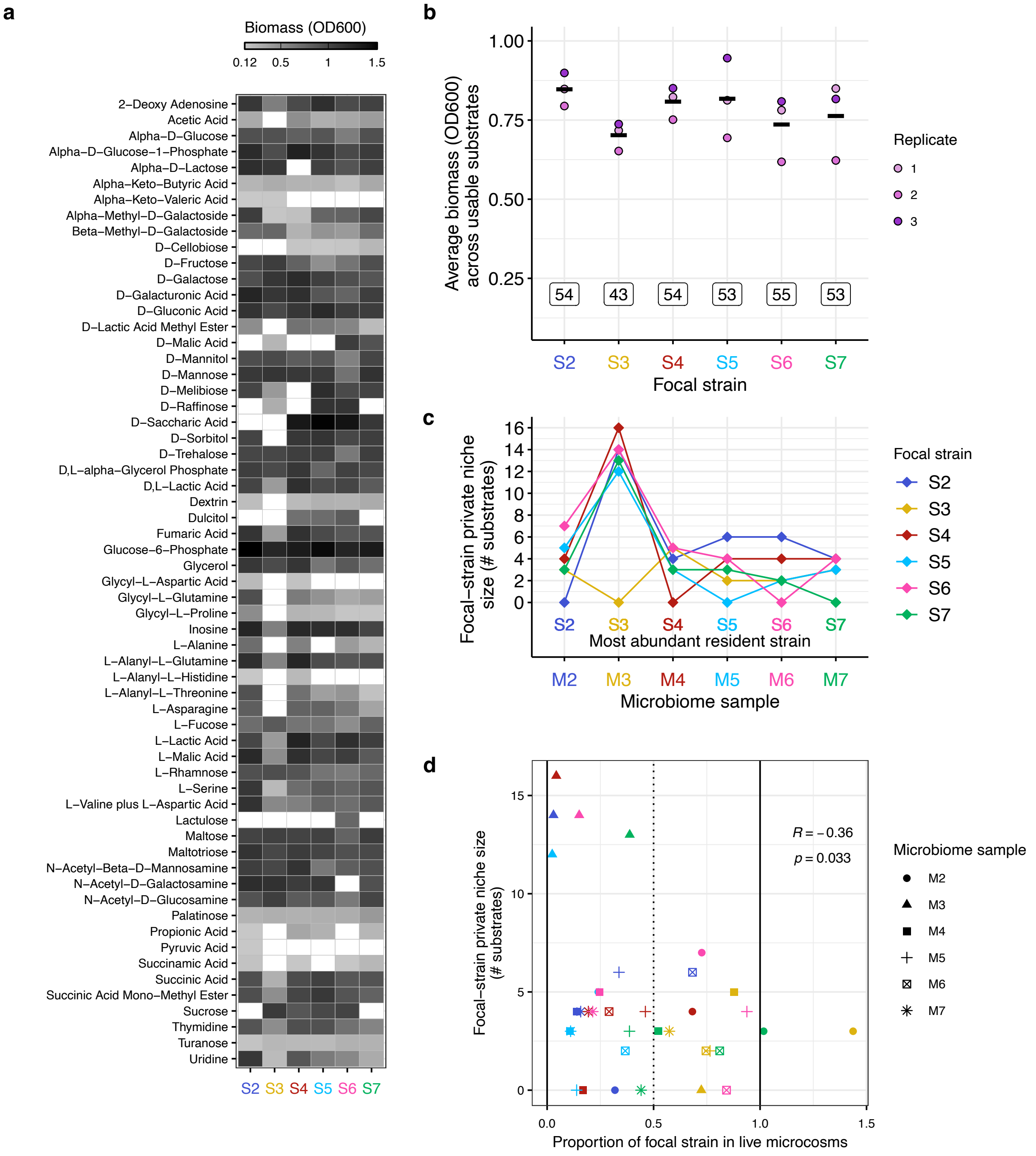


**Supplementary Fig. 7 Exploring the link between incoming strain's private niche and ecological success**

**a** Biomass produced after 24h of incubation (approximated by optical density at 600 nm, OD600) by each strain (columns) with each substrate (rows, alphabetical order). The grey shade in each cell reflects mean OD600 from three replicates (see legend on top), except in cases where biomass production was negligible (i.e., mean minus standard error <0.1, see Supplementary Methods and Supplementary Fig. 6), displayed in white. Only substrates used by at least one strain are shown. **b** Average biomass produced by the six focal strains across their respective usable substrates. Each dot correspond to one replicate and the mean is shown with a crossbar. Boxes above the x-axis give the number of usable substrates for each strain. **c** Size of the private niche (computed based on the binarized version of the data shown in panel a) for each focal strain (see legend at right) in each microbiome sample (x-axis) where it faces the most abundant resident *E. coli* strain as main competitor (labeled above the x-axis; see Supplementary Methods). The private niche captures the set of conditions where the focal strain can grow but the resident cannot, making the measure directional. For example, focal strain S3 can use three substrates that resident strain S2 cannot use, while focal strain S2 can use 14 substrates that resident strain S3 cannot use, illustrating that the value depends on which strain is focal. Note that in sympatric combinations, focal and resident strains are the same, so the size of the private niche is 0. Lines connect the outcome for each focal strain across microbiome samples. **d** Correlation between focal-strain private niche size (x-axis, data from panel c) and final proportion of focal strain relative to total *E.coli* in live microcosms (y-axis, data from insets Fig. 2). Different colors distinguish the six focal strains (as in legend from panel c), and different shapes distinguish microbiome samples, each hosting the corresponding resident strain. By definition, proportions range from 0 to 1 (black reference lines), but in two combinations a value above 1 was obtained (i.e., more CFU/mL counted on plates with antibiotics than plates without antibiotics) due to noise of the method. The dotted line shows a final proportion of 0.5, where neither the focal nor the resident strain population have an advantage relative to the other. Pearson's correlation coefficient (R) and the corresponding p-value are shown.

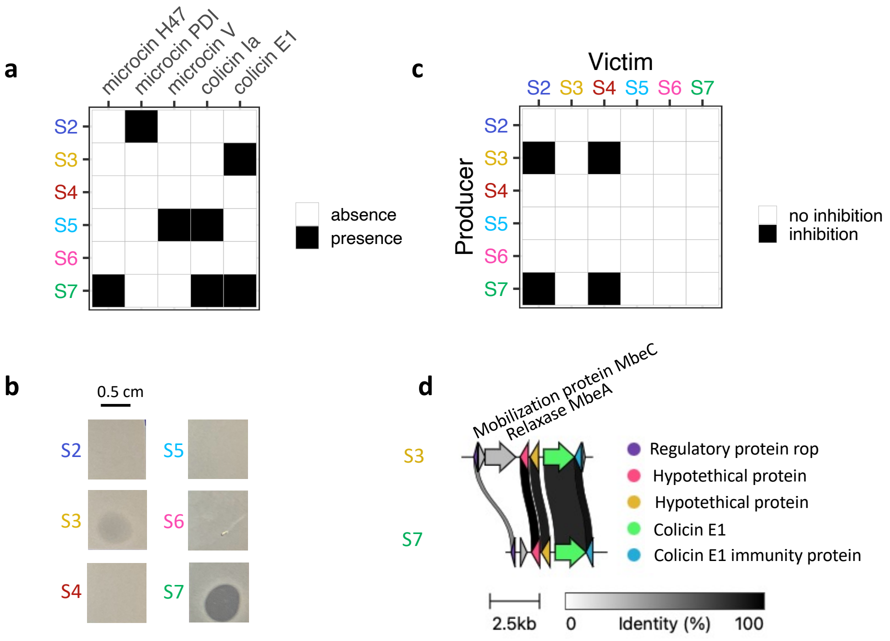


**Supplementary Fig. 8 Bacteriocin gene clusters and inhibitory phenotypes among focal strains. a** Characterized bacteriocin genes identified in the genome assemblies of the six *E. coli* strains. Putative bacteriocin-associated gene clusters were first identified with BAGEL4, and then manually curated based on their functional characterization (see Supplementary Methods). **b** Inhibition of indicator strain *E. coli* K-12 MG1655 in overlay agar by sterile filtered supernatant of stationary-phase cultures from each focal strain (labelled to the left of each subpanel). Inhibition is visible in some combinations as a clearance zone. **c** Reciprocal inhibition matrix. Here the same type of assay as in panel (b) was done with each strain-strain combination (supernatant from "Producer" strain spotted onto a lawn of "Victim" strain; inhibition recorded if there was a visible clearance zone). **d** Alignment of the colicin E1 plasmids carried by focal strains S3 and S7. Arrows represent coding sequences (CDS) . The arrows depicting CDS shared by the two plasmids are filled with matching colors and legend on the right shows their putative annotations. They are connected by vertical blocks displaying percentage identity with a white-to-black gradient (see legend on the bottom). The two CDS unique to the plasmid of S3 are labelled with putative annotations on the top. This figure was produced with clinker [12].

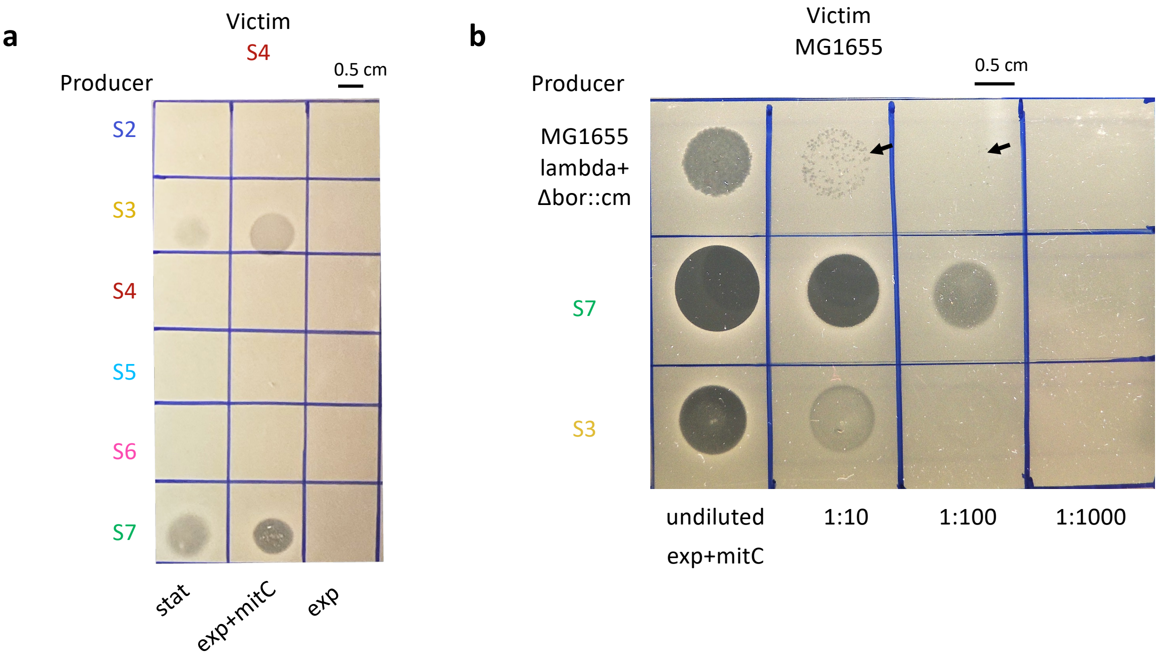


**Supplementary Fig. 9 Representative images of agar inhibition assays with different conditions or dilutions** **a** Inhibition of a focal strain in overlay agar (Victim, S4) by sterile filtered supernatant of cultures from each of the six focal strains (Producers, each row, labelled on the left) in different conditions (each column, labelled on the bottom, see abbreviations below). **b** Inhibition of indicator strain *E. coli* K-12 MG1655 in overlay agar by various dilutions (each column, labelled on the bottom) of one sterile filtered supernatant from an exponential phase culture exposed to mitomycin C of the strain K12-MG1655 lambda+ Δbor::cm [14] and the focal strains S7 and S3 (Producers, each row ,labelled on the left). Arrows show spots where plaques resulting from lysis by phages were visible.

stat= supernatant from a stationary culture; exp+mitC= supernatant from an exponential culture exposed to mitomycin C; exp=exponential culture not exposed to mitomycin C.

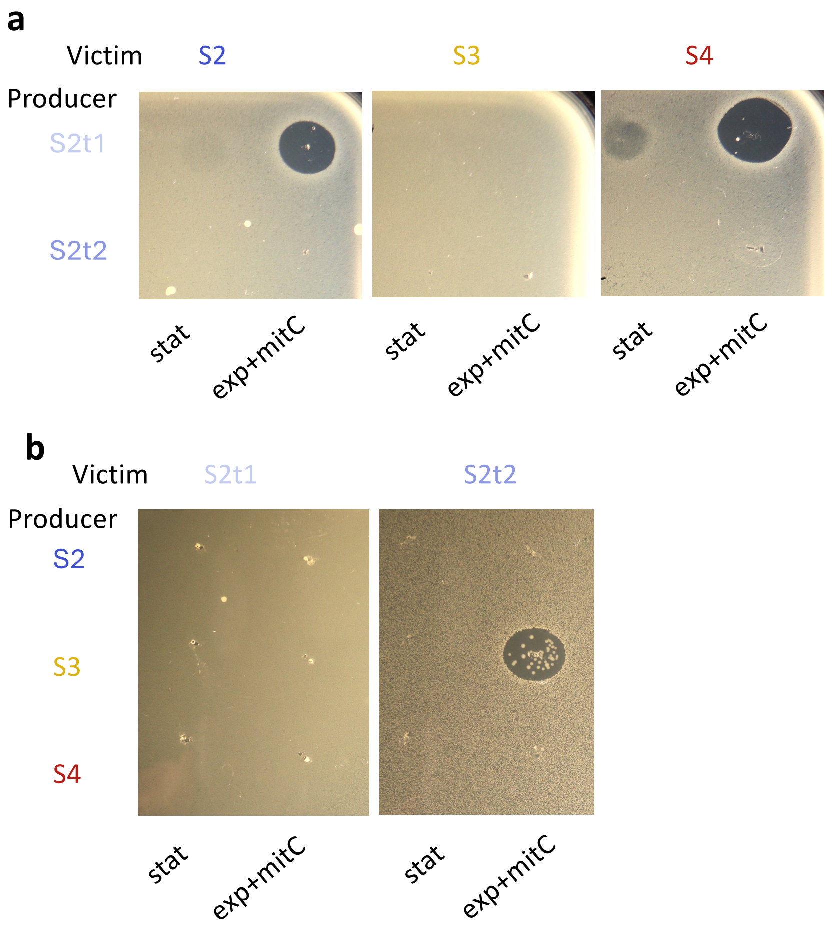


**Supplementary Fig. 10 Representative images of agar inhibition assays with colE1 transconjugants.** Transconjugants were obtained by moving the colicin E1 plasmid of S3 to S2 by agar mating, mediated by an IncF plasmid carried by S3; see Supplementary Methods). S2t1 carries both IncF plasmid and colicin E1 plasmid, S2t2 carries only IncF plasmid. **a** Inhibition of three focal strains in overlay agar (Victims, labelled on top of the three images) by sterile filtered supernatant of cultures from the two transconjugants (Producers, each row, labelled on the left) in two different conditions (each column, labelled on the bottom, see abbreviations below). **b** Inhibition of two colE1 transconjugants in overlay agar (Victims, labelled on top of the two images) by sterile filtered supernatant of cultures from three focal strains (Producers, each row, labelled on the left) in two different conditions (each column, labelled on the bottom).

Stat= supernatant from a stationary culture; exp+mitC= supernatant from an exponential culture exposed to mitomycin C.

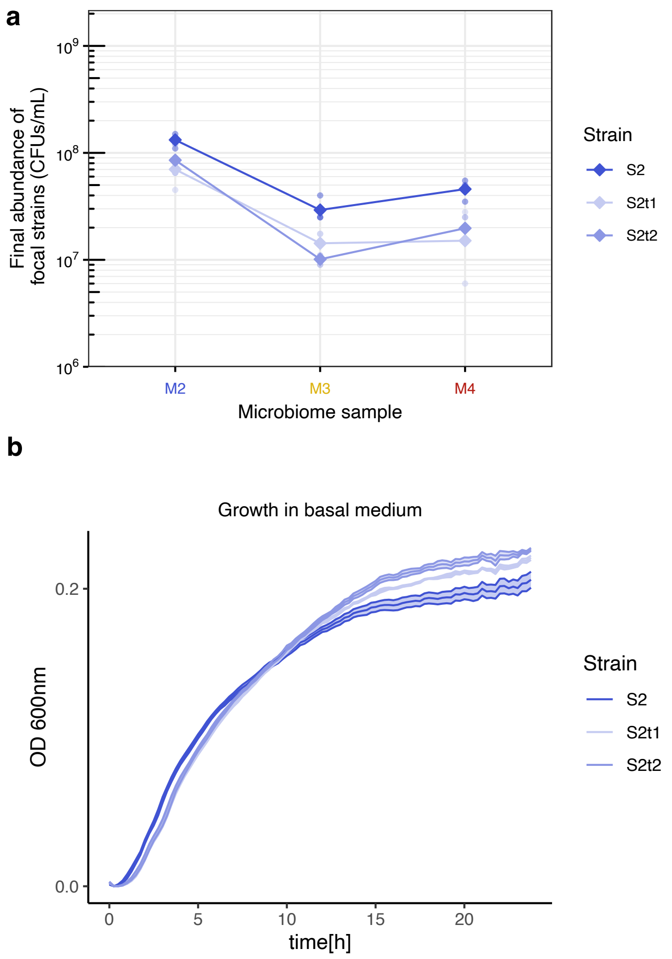


**Supplementary Fig. 11 Growth performance of ColE1 transconjugants in live microcosms a** Final abundance of focal strain S2 and transconjugants in microcosms prepared as the ones of the main experiment, with a subset of three different microbiome samples (M2-M4), inoculated with S2, S2t1 and S2t2 (total of 9 strain-microbiome combinations and thus 18 microcosms). We estimated final abundance of the focal strains by selective plating after 24h of anaerobic incubation. Diamonds represent the average (geometric mean) of three replicates for each strain-microbiome combination and dots show individual replicates. Lines connect the average for each strain across microbiome samples. Color of S2 is matched with color of its microbiome of origin (M2) and the two transconjugants are shown in clearer blue shades. S2t1 carries both IncF plasmid and colicin E1 plasmid, S2t2 carries IncF plasmid only. **b** Growth curves of S2, S2t1 and S2t2 in basal medium. The mean of three replicates is displayed with a solid line and surrounded by a shaded area representing the standard error of mean (SE).

Carrying the colE1 transconjugant did not enhance the ecological success of S2 in live microcosms; instead, both transconjugants reached consistently slightly lower final abundances than the original strain S2 **(a)**. This reduced performance likely reflects a modest fitness cost associated with plasmid acquisition, consistent with their slower growth rates in vitro **(b)**.

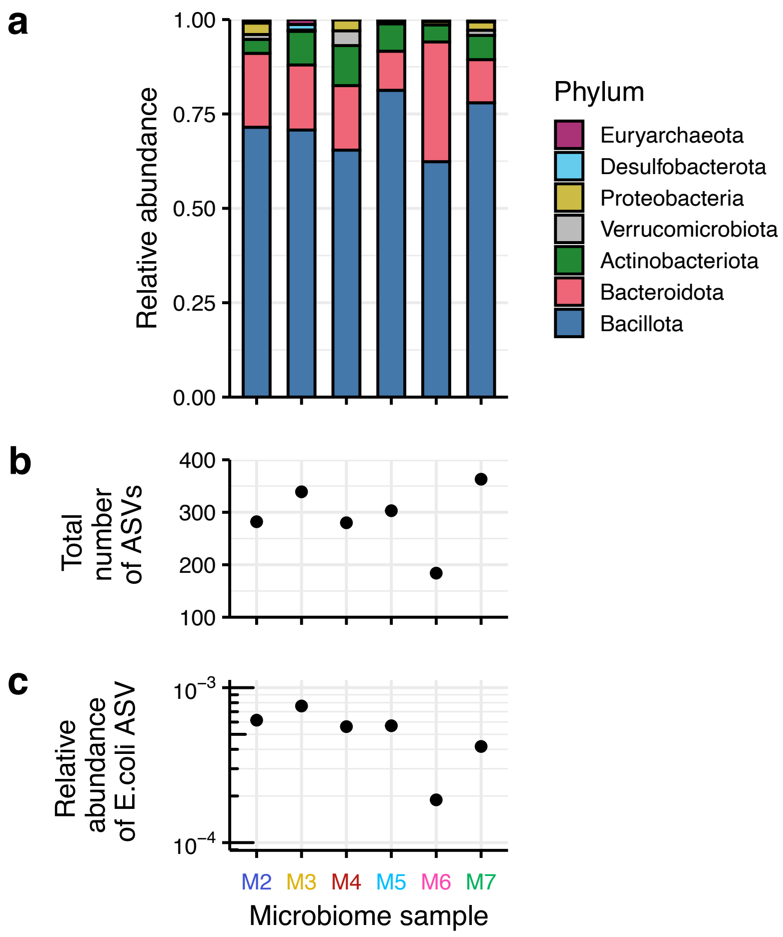


**Supplementary Fig. 12 Taxonomic composition of the six microbiome samples before cultivation. a** Relative abundance of each phylum across samples. The fourteen most abundant families belong to Actinobacteriota, Bacteroidota, Bacillota and Proteobacteria and are broken down in families in Fig. 3a **b** Total number of ASVs identified in each sample. **c** Relative abundance of the ASV assigned to *E. coli*. Because the microcosms were prepared diluting communities of ~$10^9$ CFU/mL 1:16, this range of frequencies applies to a total bacterial abundance of ~$6.25\times10^7$ CFU/mLm which corresponds to $1.2\times10^4$ CFU/mL in M6 and up to $4.8\times10^4$ CFU/mL in M3, in line with frequencies obtained based on plating (Supplementary Fig. 13a).

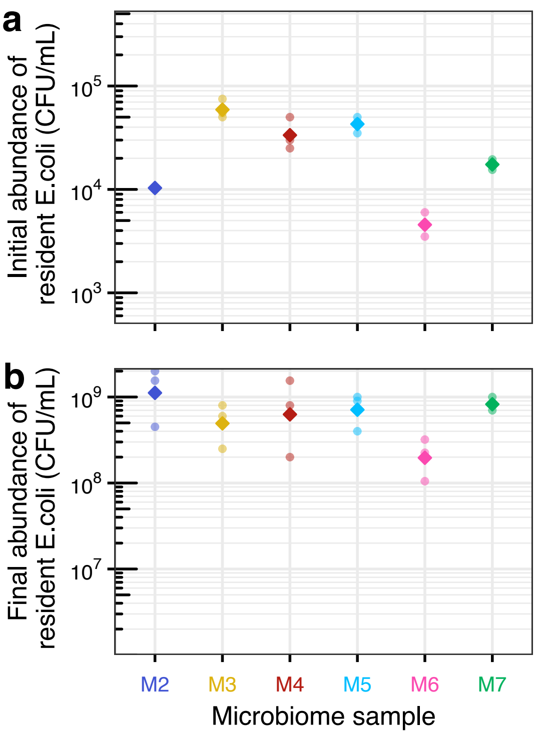


**Supplementary Fig. 13 Abundance of resident *E. coli* in live microcosms** before cultivation **(a)** and after 24h of anaerobic cultivation **(b).** We prepared these microcosms as in the main experiment but only with microbiome samples (no focal strain inoculation) and estimated resident *E. coli* abundance by plating on chromatic agar without antibiotic. Diamonds represent the average (geometric mean) of three replicates for each strain-microbiome combination and dots show individual replicates.

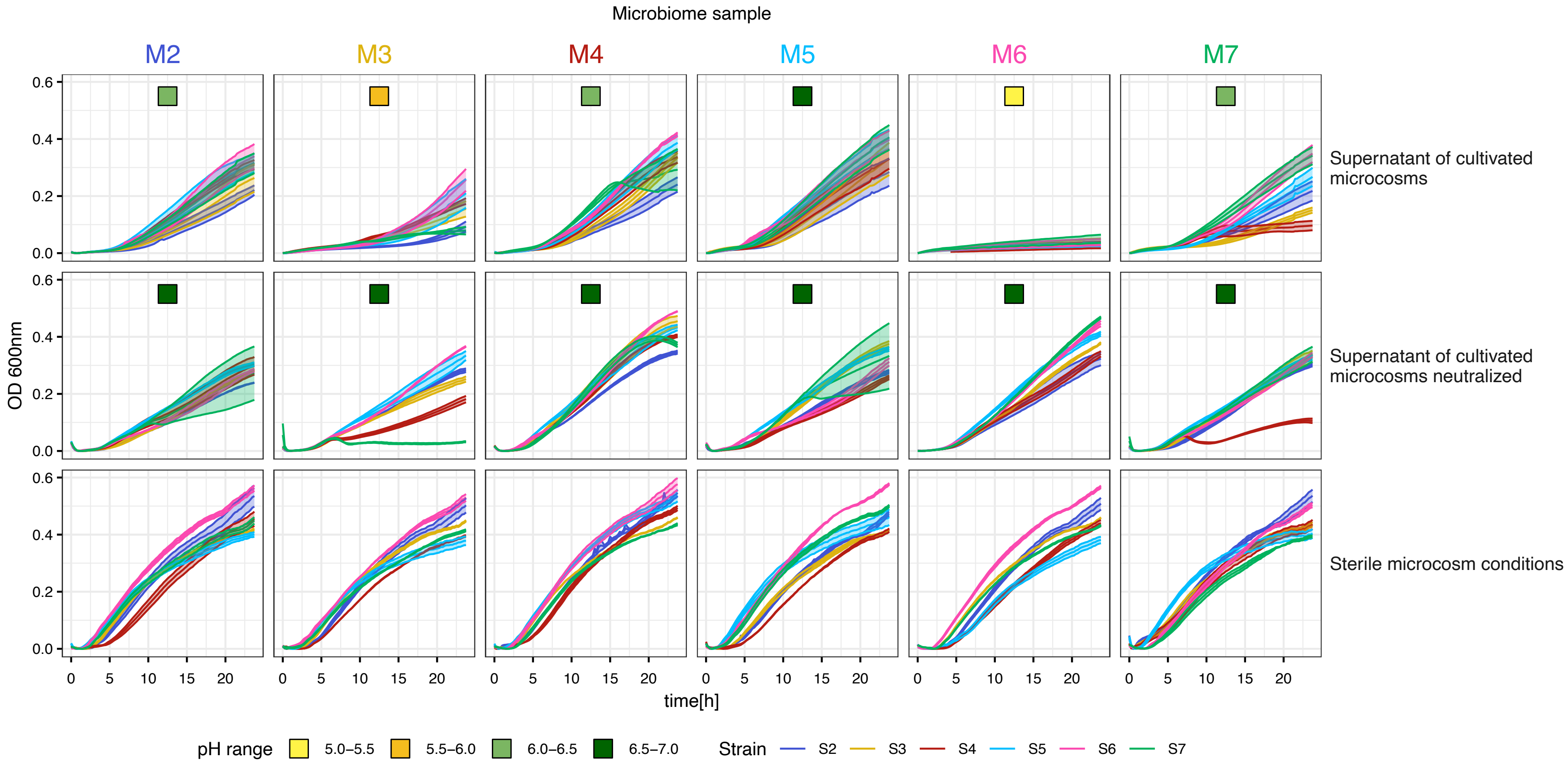


**Supplementary Fig. 14 Growth of focal strains in various conditions.** Raw data underlying Supplementary Fig. 15 (Supernatant of cultivated microcosms; unaltered or neutralized, upper and middle row respectively) and Supplementary Fig. 4b (Sterile microcosm conditions, bottom row) for reference. The colored squares on the top of the panels (for supernatant of cultivated microcosms) show approximate pH in each group on a scale from 5 to 7. The mean and standard error of mean (SE) of three replicates are displayed with a solid line surrounded by a shaded area of the same color. Data from supernatant of M6 cultivated microcosms unaltered and neutralized are also presented in Fig. 3e.

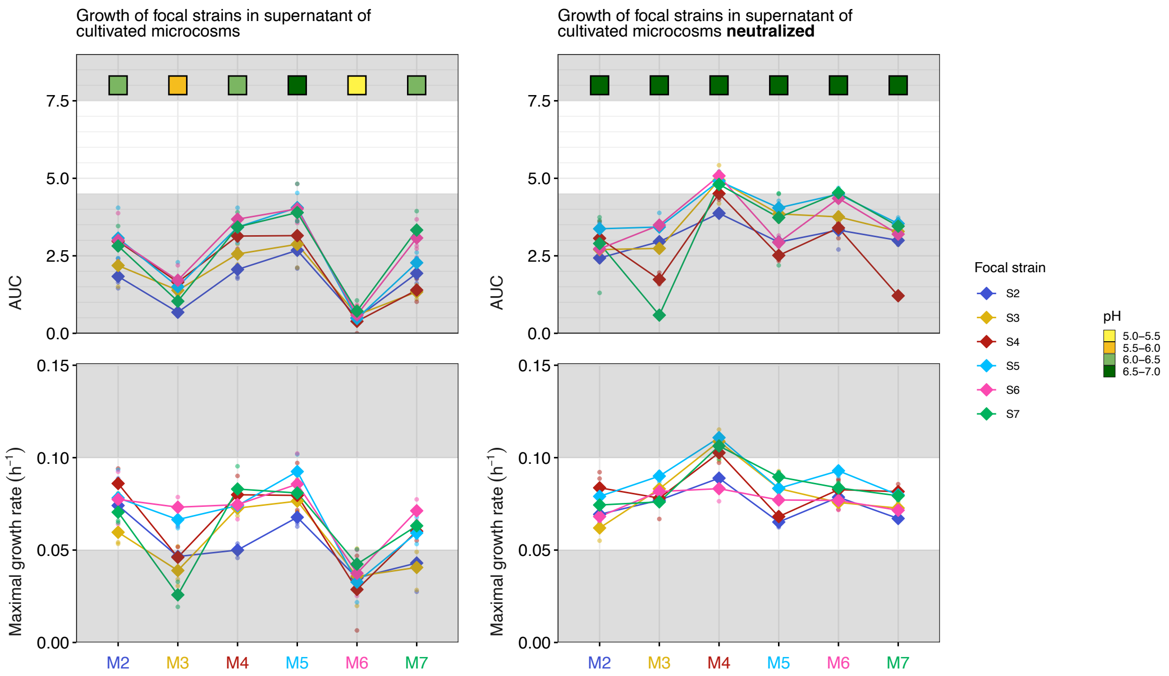


**Supplementary Fig. 15 Growth of focal strains in supernatant of cultivated microcosms**. The area under the curve (upper panels) and maximal growth rate (bottom panels) for each strain growing in supernatant of cultivated microcosms and in corresponding supernatants, neutralized to pH 6.5-7.0 (on the left and right, respectively). The raw data underlying these plots are shown in Supplementary Fig. 14. The colored squares on the top show approximate pH in each group on a scale from 5 to 7. The grayed areas serve as visual reference to compare spanned y-range across panels. The white area corresponds to the range of maximal growth rates/AUC reached by the focal stains in sterile microcosm condition (Supplementary Fig. 4b: 4.5 to 7.5 for AUC, upper panels and 0.05 to 0.10 $h^{-1}$ for maximal growth rate, bottom panels).

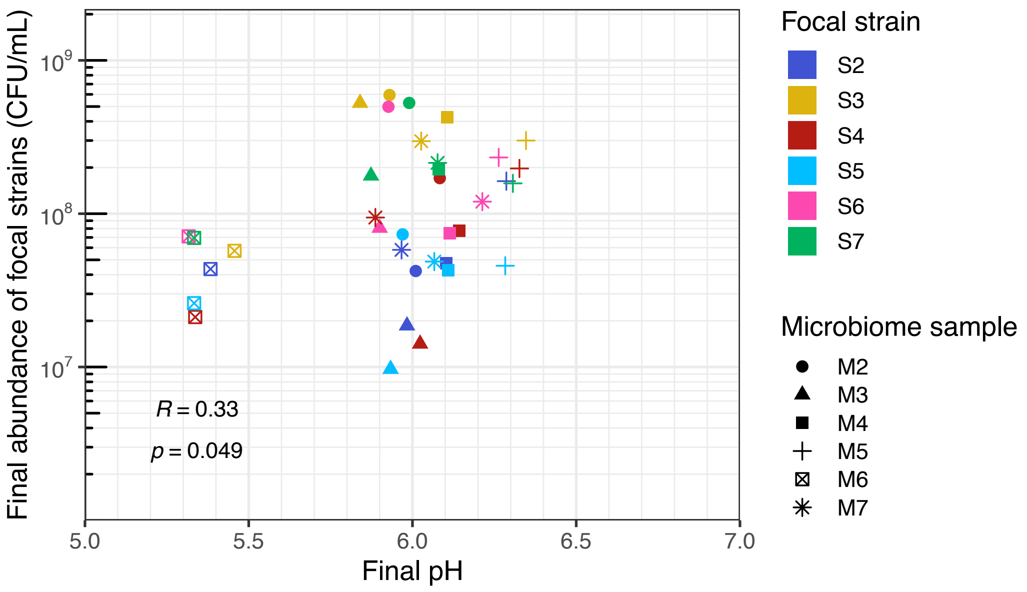


**Supplementary Fig. 16 Final pH in live microcosms and correlation with final abundance.** Correlation between final pH (Fig. 3d) and final abundance in CFU/mL (Fig. 1a). Pearson's correlation coefficient (R) and the corresponding p-value are shown. The weak correlation observed is largely driven by sample M6.

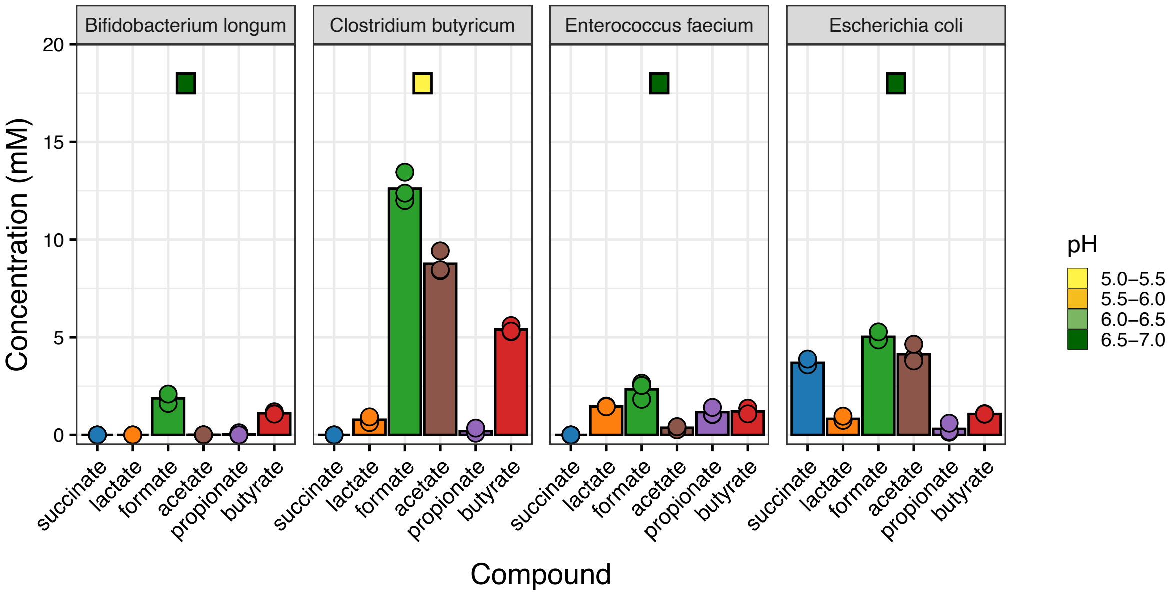


**Supplementary Fig. 17 Concentrations of fermentation products in pure cultures of four bacterial isolates.** Species of the isolate determined with MALDI-TOF are labeled on top of each panel. Bars show mean concentrations of each compound, with individual points showing replicate measurements. Compounds are labeled along the x-axis, and bars of the same compound are color-coded for clarity. Colored squares on the top of the panels indicate the final pH of the corresponding cultures, which was consistent across the three replicates.

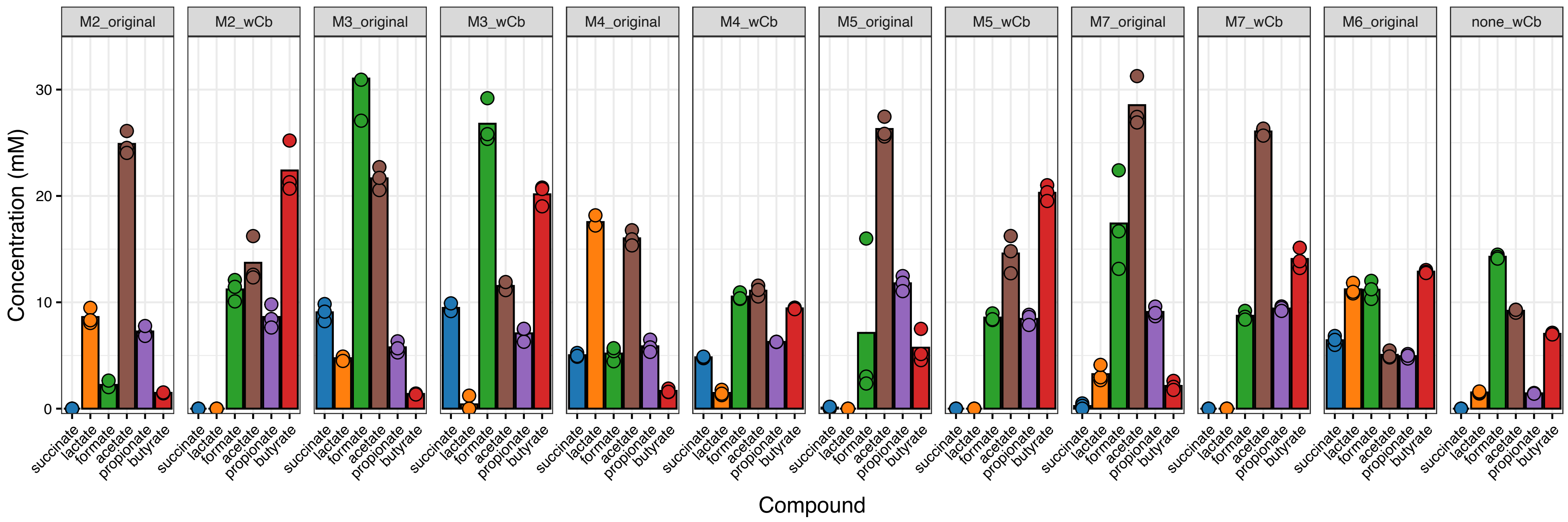


**Supplementary Fig. 18 Concentrations of fermentation products in cultivated microcosms** prepared with microbiome samples (M2, M3, M4, M5, and M7) either left untreated (*original*) or supplemented with *Clostridium butyricum* (*wCb*). Microcosm combinations (microbiome_treatment) are labeled on top of each panel. We also included triplicates of the original microbiome sample M6 (M6_original) and pure *C. butyricum* cultures (none_wCb) as a reference (on the right end). Bars show mean concentrations of each compound, with individual points showing replicate measurements. Compounds are labeled along the x-axis, and bars of the same compound are color-coded for clarity.

307

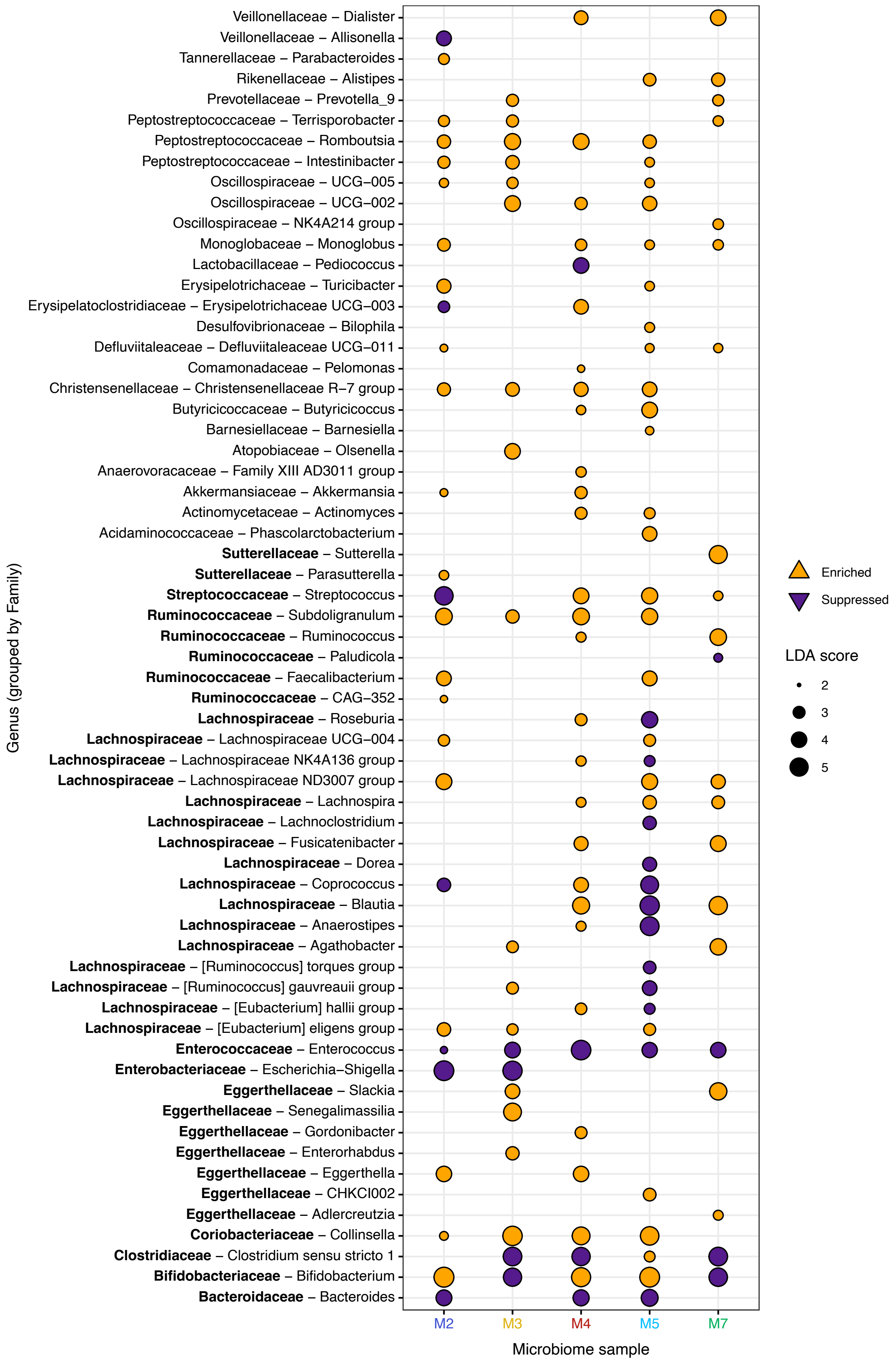

Veillonellaceae – Dialister
Veillonellaceae – Allisonella
Tannerellaceae – Parabacteroides
Rikenellaceae – Alistipes
Prevotellaceae – Prevotella_9
Peptostreptococcaceae – Terrisporobacter
Peptostreptococcaceae – Romboutsia
Peptostreptococcaceae – Intestinibacter
Oscillospiraceae – UCG–005
Oscillospiraceae – UCG–002
Oscillospiraceae – NK4A214 group
Monoglobaceae – Monoglobus
Lactobacillaceae – Pediococcus
Erysipelotrichaceae – Turicibacter
Erysipelatoclostridiaceae – Erysipelotrichaceae UCG–003
Desulfovibrionaceae – Bilophila
Defluviitaleaceae – Defluviitaleaceae UCG–011
Comamonadaceae – Pelomonas
Christensenellaceae – Christensenellaceae R–7 group
Butyricicoccaceae – Butyricicoccus
Barnesiellaceae – Barnesiella
Atopobiaceae – Olsenella
Anaerovoracaceae – Family XIII AD3011 group
Akkermansiaceae – Akkermansia
Actinomycetaceae – Actinomyces
Acidaminococcaceae – Phascolarctobacterium
Sutterellaceae – Sutterella
Sutterellaceae – Parasutterella
Streptococcaceae – Streptococcus
Ruminococcaceae – Subdoligranulum
Ruminococcaceae – Ruminococcus
Ruminococcaceae – Paludicola
Ruminococcaceae – Faecalibacterium
Ruminococcaceae – CAG–352
Lachnospiraceae – Roseburia
Lachnospiraceae – Lachnospiraceae UCG–004
Lachnospiraceae – Lachnospiraceae NK4A136 group
Lachnospiraceae – Lachnospiraceae ND3007 group
Lachnospiraceae – Lachnospira
Lachnospiraceae – Lachnoclostridium
Lachnospiraceae – Fusicatenibacter
Lachnospiraceae – Dorea
Lachnospiraceae – Coprococcus
Lachnospiraceae – Blautia
Lachnospiraceae – Anaerostipes
Lachnospiraceae – Agathobacter
Lachnospiraceae – [Ruminococcus] torques group
Lachnospiraceae – [Ruminococcus] gauvreauii group
Lachnospiraceae – [Eubacterium] hallii group
Lachnospiraceae – [Eubacterium] eligens group
Enterococcaceae – Enterococcus
Enterobacteriaceae – Escherichia–Shigella
Eggerthellaceae – Slackia
Eggerthellaceae – Senegalimassilia
Eggerthellaceae – Gordonibacter
Eggerthellaceae – Enterorhabdus
Eggerthellaceae – Eggerthella
Eggerthellaceae – CHKCI002
Eggerthellaceae – Adlercreutzia
Coriobacteriaceae – Collinsella
Clostridiaceae – Clostridium sensu stricto 1
Bifidobacteriaceae – Bifidobacterium
Bacteroidaceae – Bacteroides
Genus (grouped by Family)
M2
M3
M4
M5
M7
Microbiome sample
Enriched
Suppressed
LDA score
2
3
4
5

**Supplementary Fig. 19** LEfSe analysis performed on genus-level taxonomic composition of the cultivated gut microcosms, with reads attributed to *C. butyricum* removed prior to analyis. Analyses were performed separately for each microbiome sample, with genera aggregated from ASVs. Only genera with an LDA score >2 in at least one microbiome are shown. Each point represents a bacterial genus in a specific microbiome, with the size proportional to the absolute LDA score and color indicating direction of the effect (orange=enriched, purple=suppressed in *C. butyricum*-inoculated microcosms relative to uninoculated controls). Genera are grouped by their parent family, and families highlighted in bold correspond to those with significant family-level effects, visualized in Fig. 4f.

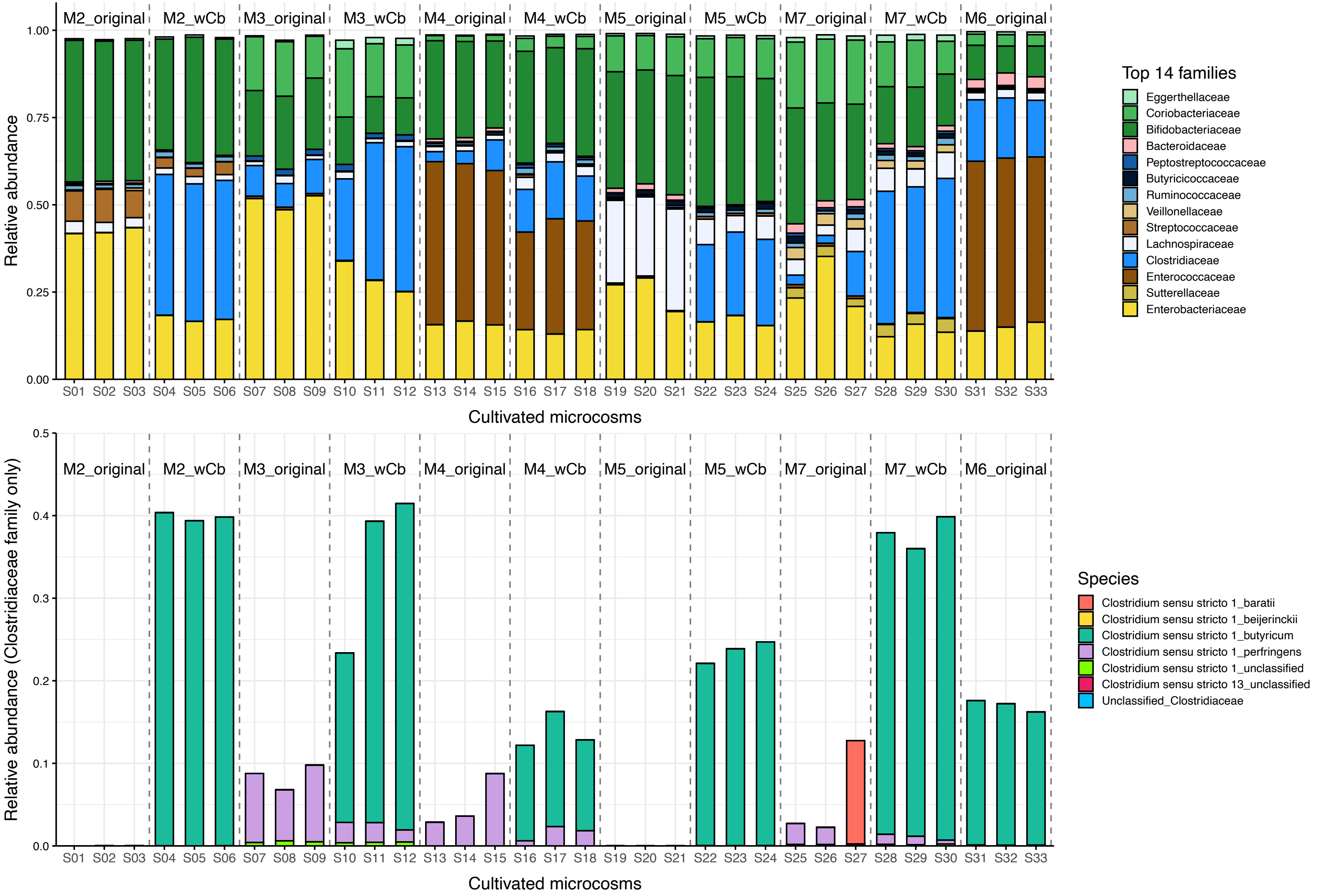
M2_original
M2_wCb
M3_original
M3_wCb
M4_original
M4_wCb
M5_original
M5_wCb
M7_original
M7_wCb
M6_original
Relative abundance
1.00
0.75
0.50
0.25
0.00
S01 S02 S03 S04 S05 S06 S07 S08 S09 S10 S11 S12 S13 S14 S15 S16 S17 S18 S19 S20 S21 S22 S23 S24 S25 S26 S27 S28 S29 S30 S31 S32 S33
Cultivated microcosms
Top 14 families
Eggerthellaceae
Coriobacteriaceae
Bifidobacteriaceae
Bacteroidaceae
Peptostreptococcaceae
Butyricicoccaceae
Ruminococcaceae
Veillonellaceae
Streptococcaceae
Lachnospiraceae
Clostridiaceae
Enterococcaceae
Sutterellaceae
Enterobacteriaceae
Relative abundance (Clostridiaceae family only)
0.5
0.4
0.3
0.2
0.1
0.0
Species
Clostridium sensu stricto 1_baratii
Clostridium sensu stricto 1_beijerinckii
Clostridium sensu stricto 1_butyricum
Clostridium sensu stricto 1_perfringens
Clostridium sensu stricto 1_unclassified
Clostridium sensu stricto 13_unclassified
Unclassified_Clostridiaceae

**Supplementary Fig. 20 Taxonomic composition of cultivated microcosms** prepared with microbiome samples (M2, M3, M4, M5, and M7) either left untreated (*original*) or supplemented with *Clostridium butyricum* (*wCb*). Labels above the plot indicate the corresponding conditions (microbiome-sample_treatment), separated by vertical dashed lines. We also included triplicates of the original microbiome sample M6 as a reference (on the right end). Sample identifiers (Sx) are indicated on the x-axis. **a** Relative abundance of the fourteen most abundant families across samples, ordered and colored based on phylum (from top to bottom: Actinobacteriota in green; Bacteroidota in pink shades; Bacillota, further distinguished by class: Negativicutes in beige shades, Bacilli in brown and Clostridia in blue shades) and Proteobacteria in yellow. Within phylum, families are ordered by ascending abundance. Enterobacteriaceae (visualized in bright yellow) made up 28.9% ± 14.4% of the cultivated microcosms where *Clostridium butyricum* had not been supplemented (including M6 microcosms) and 18.1%± 6.2% in the cultivated microcosms where it had been supplemented. **b** Relative abundance of the various species attributed to the Clostridiaceae family (visualized in vibrant sky blue above in **a**). *Clostridium butyricum* (visualized in teal) 28.9% ±11.1% of the cultivated microcosms where it had been supplemented and was absent from all where it had not been supplemented, except those prepared with its source microbiome M6, where it made up 32.2% of the community (average across replicates), and one of three M7 replicates (S26), where it occurred at a very low abundance (17/119’451 reads; 0.014%).

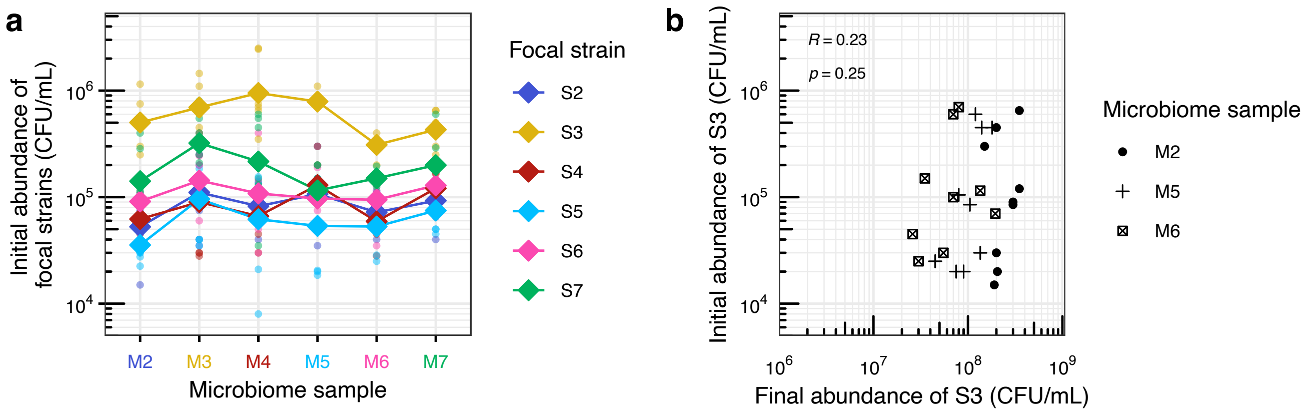


**Supplementary Fig. 21 Inoculum density and growth performance a** Initial abundance of focal strains in live gut microcosms (for which final abundance is shown on Fig. 1a). **b** Correlation between initial and final abundance of one focal strain (S3) inoculated at three different densities (three replicates per density) across microcosms prepared with three microbiome samples (M2, M5 and M6, distinguished by different shapes, legend on the right). Pearson's correlation coefficient (R) and the corresponding p-value are shown.
